# ZFP57 is a regulator of postnatal growth and life-long health

**DOI:** 10.1101/2023.08.27.554997

**Authors:** Geula Hanin, Boshra Alsulaiti, Kevin R Costello, Hugo Tavares, Nozomi Takahashi, Liudmila A Mikheeva, Anjuli Karmi Freeman, Shrina Patel, Benjamin Jenkins, Albert Koulman, Anne C Ferguson-Smith

**Affiliations:** Department of Genetics, University of Cambridge, Cambridge, CB2 3EH, UK; Core Metabolomics and Lipidomics Laboratory, Wellcome-MRC Institute of Metabolic Science, University of Cambridge, Addenbrooke’s Treatment Centre, Keith Day Road Cambridge, CB2 0QQ, UK; Wellcome-MRC Institute of Metabolic Science and Medical Research Council Metabolic Diseases Unit, University of Cambridge, Addenbrooke’s Treatment Centre, Keith Day Road Cambridge, CB2 0QQ, UK

## Abstract

Early-life factors, including nutrition, shape long-term health outcomes. Despite the essential role of lactation in maternal nutritional support, the influence of epigenetic factors on lactation and postnatal growth remains poorly understood. Zinc-finger protein 57 (ZFP57), is an epigenetic regulator of genomic imprinting, a process that directs gene expression based on parental origin, playing a vital role in mammalian prenatal growth.

Here, we identify a novel function of ZFP57 in regulating the mammary gland, where it serves as a key modulator of postnatal resource control, independently of imprinted genes. ZFP57 influences multiple aspects of mammary gland function, including ductal branching and cellular homeostasis. Its absence leads to significant differential gene expression, related to alveologenesis, lactogenesis and milk synthesis, associated with delayed lactation and altered milk composition. This results in life-long impacts on offspring including the development of metabolic syndrome.

Cross-fostering reveals intricate dynamics between mother and offspring during lactation. Pups raised by a dam of a different genotype than their birth mother exhibit exacerbated metabolic features in adulthood, providing additional insight into the programming of offspring long-term health by maternal context. This study deepens our understanding of the interplay between epigenetic factors, lactation, and postnatal resource control and identifies ZFP57 as a major regulator of both pre and postnatal resource control in mammals.

## Introduction

Genomic imprinting is a mammalian epigenetic mechanism regulating gene expression according to parental origin^1^. It is predominantly controlled by DNA methylation, influencing multiple genes. While most studies established that imprinted genes regulate prenatal growth and development, evidence suggests that imprinting is a fundamental process supporting postnatal development and maternal metabolism^2–6^.

Both the establishment and maintenance of DNA methylation at imprinted and non-imprinted regions are highly regulated^1^. One of the key regulators is ZFP57, a master regulator of genomic imprinting required for normal development^7^. Ablation of ZFP57 causes loss of DNA methylation at imprinting control regions^8,9^. In mice, deletion of the maternal gene in oocytes and the zygotic copies in early embryos causes severe loss of methylation at imprinted loci, resulting in embryonic lethality. Deletion of the zygotic ZFP57 copy causes partial neonatal lethality^7^. In humans, ZFP57 mutations have been associated with multi-locus imprinting disturbances, and transient neonatal diabetes mellitus type 1^10^.

The perinatal period is known to have major impact on the developing offspring. Various factors in early life can affect epigenetic status and gene expression, influencing life-long health^11,12^. Over the years, imprinted genes have been linked to prenatal development^1^ and nutrient transport to the embryo^4,13^. However, maternal support of the offspring continues postnatally ensuring the survival of the young, manifested as maternal care and lactation. Some imprinted genes have been associated with these processes^2,3,14^, and several are associated with adult metabolism^15–17^. Together, the perinatal and long-term effects of imprinting highlight it as a possible mechanism influencing life-long health and disease risk.

The mammary gland, a crucial organ for lactation, develops during puberty and adulthood^18^ and is a key requirement for successful milk production^19^. During pregnancy, hormonal signals drive mammary gland differentiation^20^, which yields milk-producing alveoli.

Breastmilk is essential for the early development of offspring and has been shown to influence long-term health^21^. Breastfeeding is associated with a reduced risk of obesity and diabetes, diseases which are often associated with imprinted genes^15^. Nevertheless, little is known about the role of epigenetic factors affecting lactation, postnatal growth and the developmental origins of health and disease.

## RESULTS

### ZFP57 is expressed in numerous adult somatic cells and mammary cell types

ZFP57, previously studied mainly in the embryonic context^8,22,23^ has been associated with multiple developmental processes such as cardiac development and corticogenesis^24,25^. To explore the relevance of ZFP57 in post-mitotic tissues, we compared ZFP57 expression across embryonic and adult tissues. In embryos, high expression was found in various tissues, particularly, brain regions such as the neural tube, fore-, mid- and hindbrain. Other tissues, such as the limb, heart and lung, also exhibited notable expression, while the liver showed the lowest expression of ZFP57 (Supplementary Figure 1A). Next, we compared expression levels of imprinted genes in wild-type (WT), and zygotic homozygotes (ZFP57^-/-^) which were previously described and led to ZFP57 loss-of-function^8,9^. Absence of ZFP57 in embryonic brain affected the expression of several imprinted genes at E12.5, including Zac1, Rasgrf1 and Nnat (Figure 1A). In adults, ZFP57 was highly expressed in organs such as the placenta, ovary, testis and brain regions. The lung, adipose tissue, kidney and mammary gland demonstrate lower expression of ZFP57 (Supplementary Figure 1B). Experimental validation confirmed high expression in the brain, spinal cord, and cultured embryonic stem cells (ESCs), and lower expression in adult somatic tissues including the lung and mammary gland (Supplementary Figure 1C).

**Figure 1.**
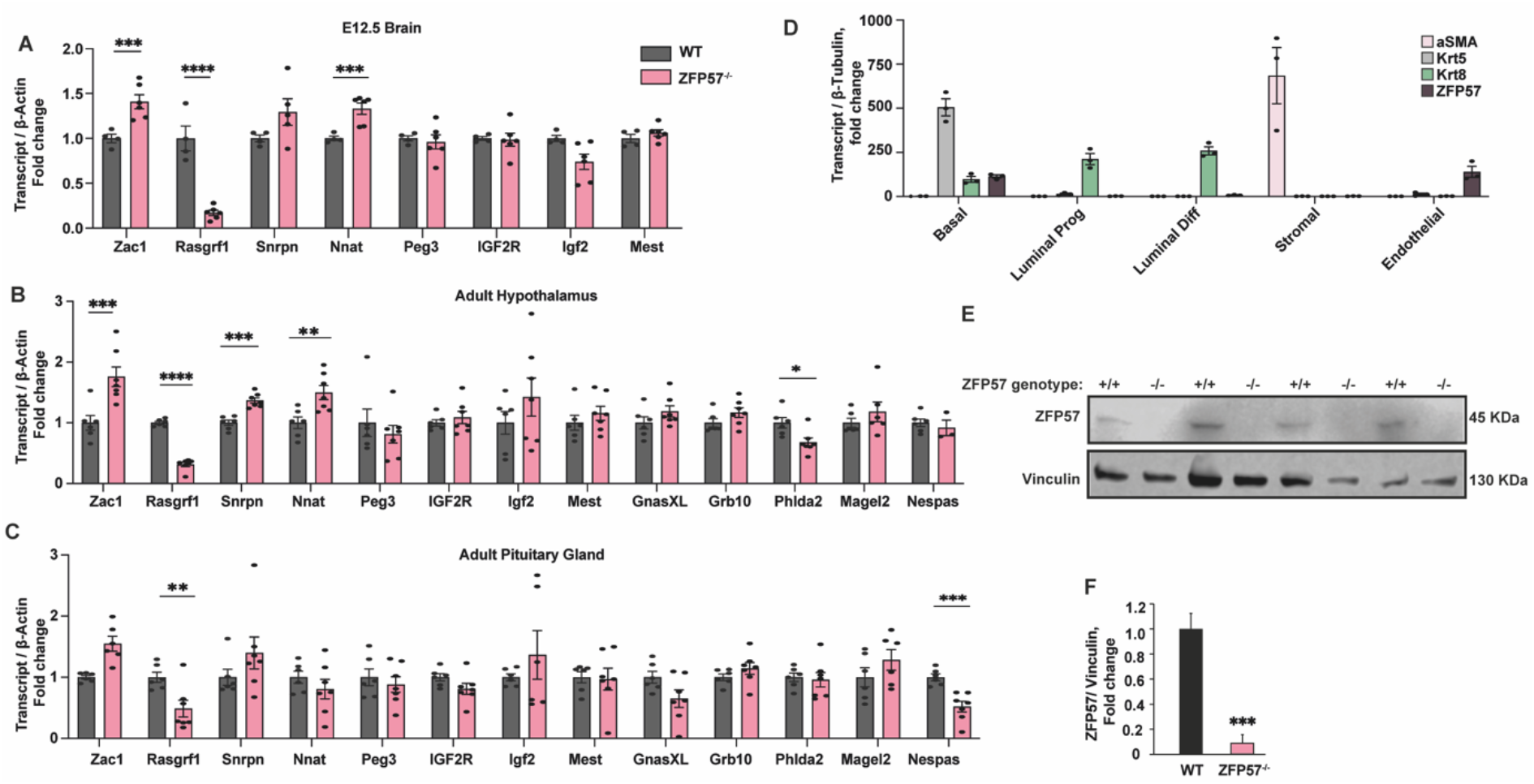
ZFP57 is relevant in somatic cells and expressed in various mammary gland cell types throughout adulthood. **(A)** Expression levels of imprinted genes associated with ZFP57 in WT and ZFP57^-/-^ E12.5 brains (*n*=4-6/time point and genotype, from 3 litters). Data are mean ± SEM, Two-tailed Student’s *t*-test. **(B-C)** Hypothalamic **(B)** and pituitary gland **(C)** expression levels of imprinted genes associated with ZFP57 or maternal behaviour and milk let-down in WT and ZFP57^-/-^ mice (*n*=6-7/time point and genotype) Data are mean ± SEM, Two-tailed Student’s *t*-test. **(D)** ZFP57, αSMA, Krt5 and Krt8 expression levels in sorted mammary cell types from WT mice at lactation day 2 (*n*=3/genotype). Data are mean ± SEM normalised to β-Tubulin. **(D-E)** Western blot **(D)** and quantification **(E)** of ZFP57 protein levels normalised to Vinculin in ZFP57^-/-^ and WT mammary glands at lactation day 2 (*n*=4 mice per genotype). Data are mean ± SEM, Two-tailed Student’s *t*-test. \**P* < 0.05, \*\**P* < 0.01, \*\*\**P* < 0.001, \*\*\*\**P* < 0.0001;

ZFP57 is highly expressed in the placenta, which is crucial for resource allocation during prenatal stages. The placenta and mammary gland share similar functions, supporting offspring growth through nutritional resource control^6^. This led us to hypothesize that ZFP57 evolved as an upstream regulator supporting offspring growth both pre- and postnatally Maternal behaviour can impact lactation performance and involves imprinted gene function in the hypothalamus and pituitary gland^15,27^ where ZFP57 levels are high. To initiate an exploration of the role of ZFP57 in postnatal resource control, we quantified the expression of imprinted genes regulated by ZFP57^8^ in adult mice hypothalami and pituitary glands. In the hypothalamus, we observed changes in several imprinted genes regulated by ZFP57, including Zac1, Rasgrf1, Snrpn and Nnat, and a reduction in Phlda2 levels which may affect maternal care^28^ (Figure 1B). In the pituitary gland, both Rasgrf1 and Nespas showed a similar pattern (Figure 1C). This indicates that imprinted gene regulation by ZFP57 is relevant in both embryonic and adult tissues.

To investigate ZFP57 expression in the mammary gland throughout its development, we quantified ZFP57 expression in five mammary cell populations from WT female mice, on lactation day 2. We sorted basal cells (Lin^-^CD31^neg^CD45^neg^EpCAM^lo^ CD49f^hi^), luminal differentiated cells (Lin^-^CD31^neg^CD45^neg^EpCAM^high^ CD49f ^low^ CD49b ^low^), luminal progenitor cells (Lin^-^CD31^neg^CD45^neg^EpCAM^high^ CD49f ^low^ CD49b ^high^), immune cells (CD45^+^) and endothelial cells (CD31^+^). Supplementary Figure 2A provides an overview of the gating strategy. ZFP57 was expressed in basal and endothelial cells. However, the level of ZFP57 expression compared to the basal luminal and stromal markers Krt5, Krt8 and αSMA was lower. Notably, luminal differentiated, luminal progenitors and stromal cells exhibited the lowest expression (Figure 1D). These findings suggest that ZFP57 may function specifically in basal and endothelial cells during the adult mammary gland developmental cycle.

To evaluate the presence of ZFP57 protein in mammary glands, we extracted protein from lactating mammary glands of both WT and ZFP57^-/-^ mice. Protein blot quantifications revealed the presence of the protein in WT mice albeit at low levels, and the absence in ZFP57^-/-^ mice (Figure 1E-F).

### ZFP57 affects mammary ductal branching during the reproductive cycle

To unravel the role of ZFP57 in the mammary gland, we focused on key developmental stages during which the gland prepares to support offspring growth. We examined the architecture of nulliparous mammary glands, gestation days 4.5, 9.5, and 14.5, and lactation day 2. Comparing WT and ZFP57^-/-^ mammary glands, revealed aberrant tertiary branching in ZFP57^-/-^ glands (Figure 2A). Nulliparous ZFP57^-/-^ mammary glands showed an increased number of ducts and total branching points compared to WT controls, reminiscent of early pregnancy glands in WT animals. In contrast, in gestation days 4.5, 9.5 and 14.5, ZFP57^-/-^ mice showed a significant reduction in ducts and branching points (Figure 2A-B). This indicates that ZFP57 contributes to normal mammary tertiary branching and that its absence leads to precocious development, potentially impacting tissue functionality.

**Figure 2.**
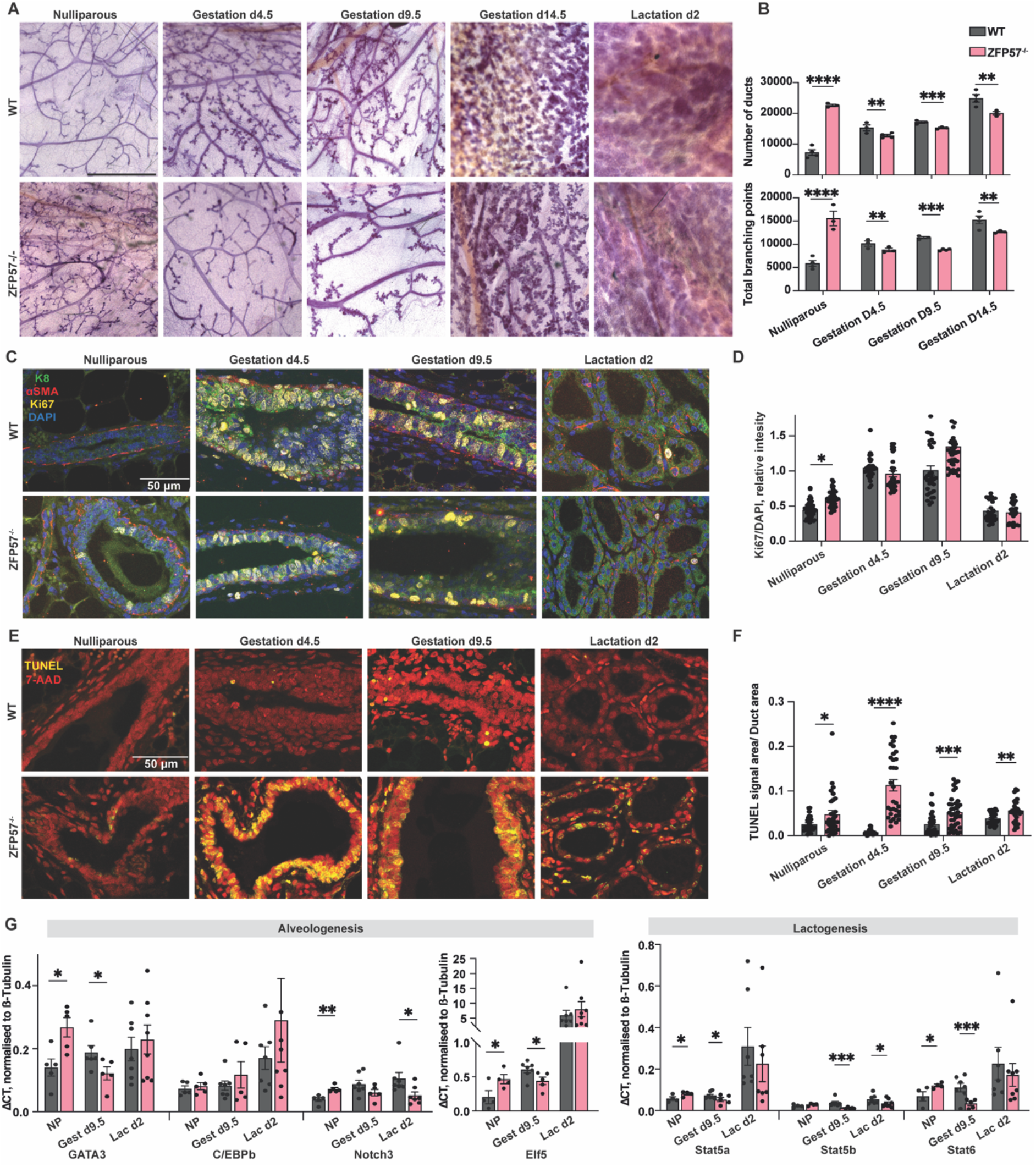
ZFP57^-/-^ mice exhibit abnormal mammary gland morphology and cellular dynamics. **(A)** Representative mammary gland whole mounts stained with Carmine Alum, from ZFP57^-/-^ and WT mice at nulliparous, gestation day 4.5, 9.5 and 14.5 and lactation day 2. Size bar represents 20 mm. **(B)** Quantified number of ducts and total branching points of mammary gland whole mounts. (*n*=3-4/stage and genotype). Data are mean ± SEM. One-way ANOVA with Tukey’s corrections. **(C-D)** Representative immunofluorescence **(C)** and quantification **(D)** of Ki67 in sectioned mammary glands from ZFP57^-/-^ mice compared with WT across nulliparous, gestation day 9.5 and lactation day 2 stages. Krt8 and αSMA served as luminal and basal markers respectively. DAPI served as a nuclear marker. *n*=3/genotype and stage, 10 fields quantified/section. Data are mean ± SEM. One-way ANOVA with Tukey’s corrections. **(E-F)** Representative images **(E)** and quantification **(F)** of apoptotic nuclei in ZFP57^-/-^ and WT mammary glands using TUNEL staining. 7-AAD was used as a nuclear marker. *n*=3-5/genotype and stage, 10 fields quantified/section. Data are mean ± SEM. Two-way ANOVA with Tukey’s corrections. **(G)** Relative alveologenesis (left) and lactogenesis (right) related transcripts in whole mammary tissues from ZFP57^-/-^ mice compared with WT across nulliparous, gestation day 9.5 and lactation day 2 stages. *n*=4-5. Values are normalised to β-Tubulin. Data are mean ± SEM. Two-way ANOVA with Tukey’s corrections. * p<0.05, ** p<0.01, *** p<0.001. *** p<0.001, **** p<0.0001.

Interestingly, on lactation day 2 the gross appearance of the mammary gland seemed normal and fully populated with alveoli, suggesting a potential compensation and ability to produce milk.

### ZFP57 regulated mammary cellular dynamics in-vivo

Next we tested the physiological functionality of gland. The healthy development of mammary glands relies on homeostasis between mammary cellular populations^29^. To investigate whether the aberrant mammary branching in ZFP57^-/-^ mice relates to abnormal homeostasis, we used flow cytometry to compare cellular proportions in ZFP57^-/-^ and WT mammary glands. Loss of ZFP57 resulted in a significantly imbalanced cellular composition throughout the mammary gland developmental cycle. Specifically, ZFP57^-/-^ glands had altered proportions of epithelial basal and luminal cells in nulliparous glands, at gestation day 9.5 and lactation day 2. Further quantification of luminal subpopulations revealed a higher fraction of differentiated luminal cells during gestation. During lactation a significant proportion of them retained their progenitor state, potentially affecting tissue functionality. (Supplementary Figure 3A). Endothelial cells were decreased in ZFP57^-/-^ glands compared to WTs in all stages (Supplementary Figure 3B). The absolute numbers of sorted cells and the changes per stage and genotype are presented (Supplementary Figure 3C-E). These findings suggest that ZFP57 loss results in an imbalanced mammary cellular composition, particularly in mammary epithelial cells.

Given the imbalance between proliferative and differentiated epithelial cells, we immunostained mammary sections for the Ki67 proliferative marker. Quantification of Ki67-positive cells showed that ZFP57^-/-^ nulliparous glands exhibited positive Ki67 cells in contrast to WT controls, consistent with the histological analysis. During gestation days 4.5, 9.5 and lactation day 2, no significant differences were observed (Figure 2C-D), further indicating premature proliferation in ZFP57^-/-^ nulliparous glands.

Together, the data suggest that in mutants, premature ductal proliferation occurs in nulliparous glands, followed by a significant decrease in ducts and branching points on gestation day 4.5. To explore whether this reduction is reflected at the molecular level, we performed TUNEL staining, which revealed a significantly higher fraction of apoptotic mammary cells in ZFP57^-/-^ compared to WT sections, starting from gestation day 4.5. This pattern persists at gestation day 9.5 and lactation day 2. (Figure 2E-F). Quantitative real-time PCR analysis of differentiation-related genes in whole mammary tissue confirmed that ZFP57^-/-^ mice experience premature alveologenesis and lactogenesis in nulliparous glands, indicated by increased levels of GATA3, Notch3, Elf5, Stat5a, Stat6; while WT mammary glands typically exhibit an increase in these transcripts as they undergo alveologenesis and lactogenesis^30^. This was followed by a significant decrease in the levels of GATA3, Elf5, Stat5a, Stat5b, and Stat6 during gestation, in contrast to the expected increase in these transcripts during pregnancy. During lactation, most of these transcripts remain unchanged, except for Notch3 and Stat5b which remain lower compared to WT glands (Figure 2G).

Together, our data shows that ZFP57^-/-^ results in precocious development of the nulliparous mammary gland characterised by extensive tertiary branching, tissue proliferation and upregulated lactogenesis and alveologenesis-related genes. However, during gestation, ZFP57^-/-^ glands experience a significant decrease in mammary ducts and branching, along with extensive apoptosis and reduced activation of the transcriptional network that normally transforms the mammary gland into a physiologically-functional organ during lactation.

### ZFP57 does not act through imprinting in the mammary gland but affects mammary gland development and milk synthesis genes

To elucidate the transcriptional changes contributing to mammary gland impaired homeostasis, we analysed transcriptomes from 120 sorted cell populations at various stages: nulliparous, gestation day 9.5, and lactation day 2. Characterisation of transcriptome-wide changes using principal component analysis (PCA) revealed that variation of the samples is primarily affected by cell type and stage but not genotype (Figure 3A-B), indicating a lack of major transcriptional differences in ZFP57^-/-^ glands.

**Figure 3.**
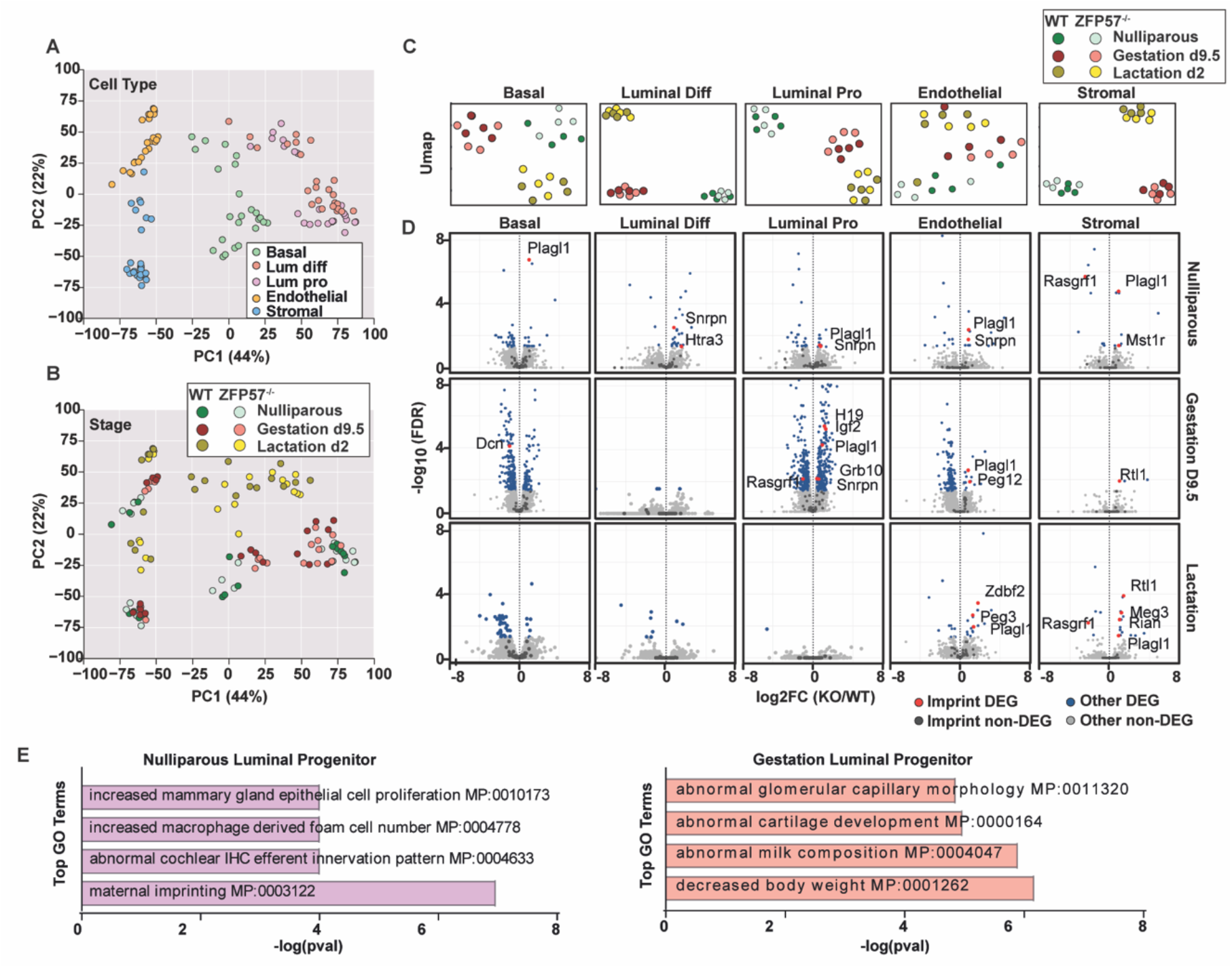
Epigenetic and transcriptional profiling of mammary tissue in ZFP57 ^-/-^ mice. **(A-B**) Principal component analysis (PCA) of mammary gland gene expression comparing ZFP57^-/-^ and WT mice. Colours represent different purified cell-types **(A)** and developmental stage separated by genotype **(B).(C)** UMAP visualisation of cell-type specific RNA-seq libraries, coloured by stage and genotype **(D)** Volcano plot representing differentially-expressed imprinted and non-imprinted genes in ZFP57^-/-^ compared WT mice across 5 different cell populations and 3 developmental stages. **(E)** Pathway enrichment analysis of biological processes that are changed in ZFP57^-/-^ nulliparous luminal progenitor cells, and during gestation. The top 4 GO terms are shown, arranged by their log(p-value). *n*=4 mice/genotype cell type and stage.

Furthermore, cell-type-specific clustering of these transcriptomes showed that ZFP57^-/-^ and WT maintain their clustering by stage. However, in luminal progenitors, and endothelial cells, genotypic separation occurs during gestation. In stromal cells, separation between genotypes was observed during lactation (Figure 3C).

To study whether ZFP57^-/-^ is associated with impaired genomic imprinting in the mammary gland, we plotted differentially-expressed genes (DEGs) in these libraries. Only 15 imprinted genes were differentially expressed in ZFP57^-/-^ glands. Of those, none of the imprinted genes within the same imprinted control region showed the reciprocal behaviour expected if imprinting were lost in the absence of ZFP57 (Figure 3D, Supplementary Figure 4). This indicates that perturbed expression of imprinted genes is at the level of transcription rather than through loss of ZFP57-mediated imprinting control.

Importantly, most of the DEGs were observed in luminal progenitor cells (Figure 3D), which exhibit minimal expression of ZFP57 (Figure 1C). This suggests that the observed phenotype in the mammary gland could occur as a downstream effect of loss of ZFP57 function either earlier in development or elsewhere. However, analysing the enriched pathways among the DEGs in ZFP57^-/-^ luminal progenitors, revealed several terms related to lactation.

Specifically, in the nulliparous stage, among the top 4 GO terms was mammary gland epithelial proliferation. Similarly, during gestation d9.5, the top 4 terms included milk composition and body weight (Figure 3E). Together, this implies that ZFP57 plays a role in the regulation of genes essential for normal mammary gland development and milk production, that this occurs independently of imprinted genes in the mammary gland, and that this may be caused by functions of ZFP57 earlier development and/or at distant sites regulating mammary gland physiology such as the brain. Further extensive in vivo analysis is required to elucidate these possibilities.

### ZFP57^-/-^ dams show delayed lactation onset and altered milk composition

To examine how the observed transcriptional and morphological aberrations observed in ZFP57^-/-^ mammary glands impact lactation and maternal nutritional resources, we focused on two prominent mouse crosses to assess maternal effects of ZFP57 mutants on offspring wellbeing: WT females crossed to a ZFP57^-/-^ males, generating control paternal heterozygous pups (ZFP57^+/-p^), or ZFP57^-/-^ females crossed to WT males generating maternal heterozygous pups (ZFP57^m-/+^). Both crosses yield genetically identical heterozygous offspring of a single genotype, which do not exhibit loss of imprinting^8^. ZFP57^-/-^ females did not display any differences in latency to get pregnant (Supplementary Figure 5A), suggesting no major hormonal imbalance. This was supported by unchanged levels of circulating progesterone and oestradiol in ZFP57^-/-^ nulliparous females compared to WTs (Supplementary Figure 5B-C). ZFP57^m-/+^ pups born to ZFP57^-/-^ dams showed a slight increase in birth weight (Figure 4A). This could potentially be attributed to longer gestation period in ZFP57^-/-^ dams compared to WTs (Figure 4B). We observed that ZFP57^-/-^ generally had smaller litters compared to WT dams (Figure 4C). Notably, pups born to ZFP57^-/-^ dams had a lower early postnatal survival rate during the lactation period compared to those born to WT or ZFP57^+/-^ dams (Figure 4D). This could be explained by a delay in milk secretion as evidenced by delayed appearance of milk spots in pups born to ZFP57^-/-^ dams (Figure 4E).

**Figure 4.**
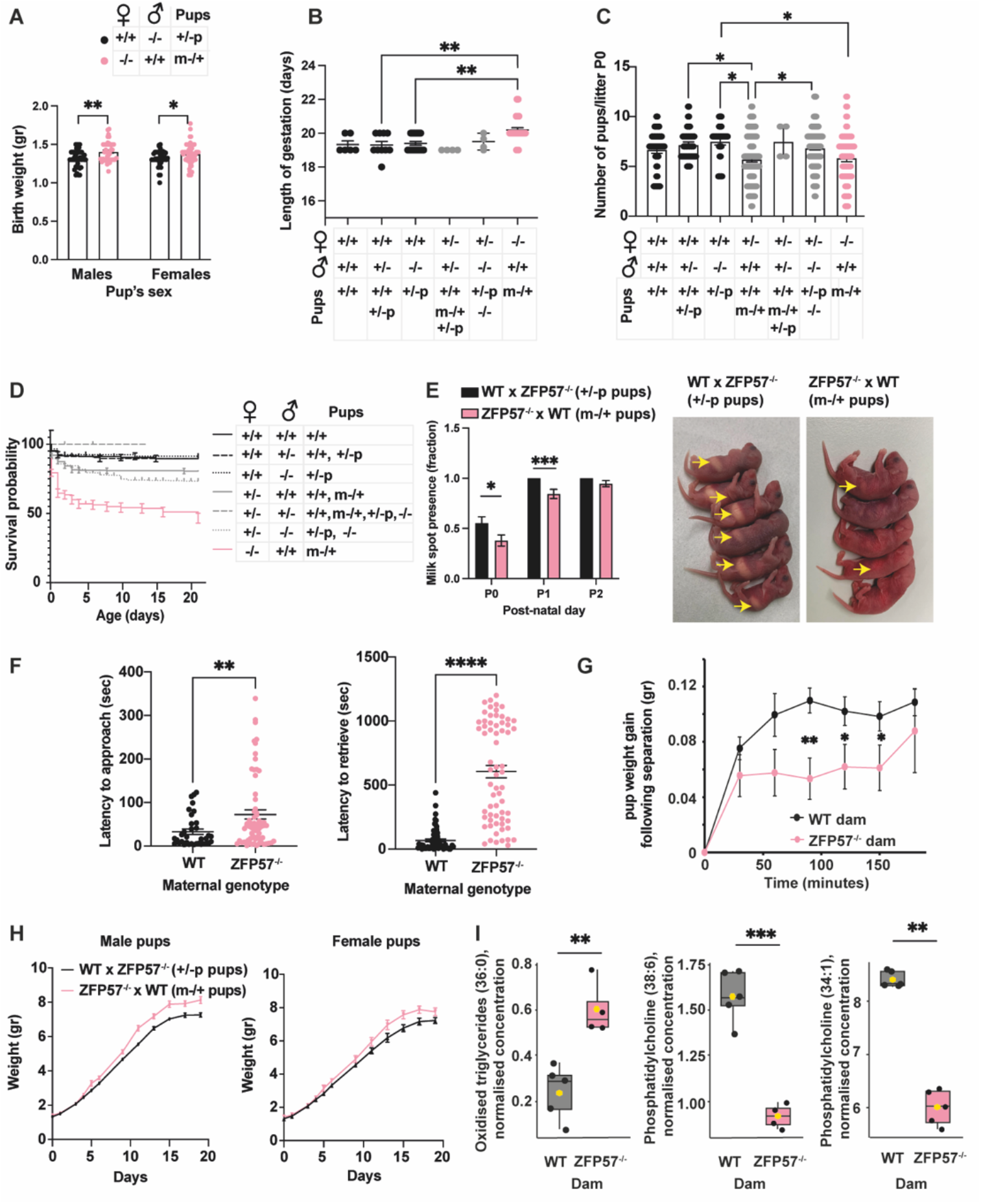
Loss of ZFP57 results in delayed lactation onset, altered milk composition and excessive weight gain. **(A)** Birth weight of heterozygous pups born to ZFP57^-/-^ or WT dam. *n*=36-53, 13–16 litters/genotype. Data are mean ± SEM. One-way ANOVA with Tukey’s corrections. **(B)** Length of gestation for each cross. *n*=6 for WTxWT and ZFP57^-/-^xZFP57^+/-^, *n*=10 for WTxZFP57^+/-^, *n*=20 for WTxZFP57^-/-^, *n*=42 for ZFP57^-/-^xWT. Data are mean ± SEM. One-way ANOVA with Tukey’s corrections. **(C)** Number of pups per litter at postnatal day 0. *n*=21-28 for WTxWT, WTxZFP57^+/-^ and WTxZFP57^+/-^. *n*=43 for ZFP57^-/-^xWT, *n*=71-83 for ZFP57^+/-^ xZFP57^-/-^ and ZFP57^+/-^xWT. Data are mean ± SEM. One-way ANOVA with Tukey’s corrections. **(D)** Kaplan–Meier curve of pups from different crosses, throughout lactation period. WTxWT, WTxZFP57^+/-^, WTxZFP57^-/-^ *n*= 158-201, 21-28 litters, ZFP57^+/-^xWT, ZFP57^-/-^xWT *n*= 407-569, 71-83 litters, ZFP57^+/-^xZFP57^+/-^ *n*=15, 2 litters. Log-rank test *P* < 0.0001 for ZFP57^-/-^ xWT. **(E)** Quantification and representative pictures of pups with milk spots at postnatal days 0, 1, 2. *n*=80-94 pups from 13 litters/cross. Values are expressed as mean ±SEM. One-way ANOVA with Tukey’s corrections. **(F)** Latency of ZFP57^-/-^ and WT dam to approach and retrieve their pups. *n*=64-65 pups from 9-10 litters. Values are mean ±SEM Two-tailed Student’s *t*-test. **(G)** Pup weight gain following separation (time 0), and reunion with the dam. *n*=20-32 pups from 5-6 litters. Values are mean ±SEM. Log-rank test. **(H)** Growth trajectories of pups born to ZFP57^-/-^ or WT dams, during lactation period. *n*=46-53 pups from 11-12 litters/cross. **(I)** Oxidised triglycerides (36:0) and phosphatidylcholine (38:6 and 34:1) in milk collected from ZFP57^-/-^ and WT dams on lactation day 9. *n*=4-5 dams. P < 0.05 values are mean relative concentrations (µM) ±SEM. * p<0.05, ** p<0.01, *** p<0.001. *** p<0.001, **** p<0.0001.

Additionally, impaired maternal behaviour, indicated by the pup retrieval assay, may contribute to decreased pup survival (Figure 4F). Nevertheless, our analysis did not reveal any significant difference in pup contact, licking, or grouping within the nest (Supplementary Figure 5E-G), except for a slight decrease in nest organisation at P0 that was not evident later (Supplementary Figure 5D). Moreover, we found a minor decrease in pup body temperature at P1 and P2 which is unlikely to cause hypothermia (Supplementary Figure 5H). Taken together, these findings suggest that the absence of maternal ZFP57 is associated with delayed onset of lactation and increased pup mortality.

Next, to assess milk let-down, we conducted a suckling assay and monitored the weight gain of the pups following separation from the dam. We found that ZFP57^m-/+^ pups had delayed weight recovery when reunited with their ZFP57^-/-^ mothers compared to ZFP57^+/-p^ pups with their WT mothers. However, it is worth noting that both groups eventually achieved similar weight gain. (Figure 4G). We did not observe any impaired suckling, pups were found in proximity to the nipple and often latched on during weighing. This suggests the possibility of either compromised milk let-down by ZFP57^-/-^ dams, or a suckling issue specific to ZFP57^m-/+^ pups.

To evaluate whether this observation persists throughout lactation, we monitored the growth of ZFP57^m-/+^ and ZFP57^+/-^ pups raised by ZFP57^-/-^ and WT mothers respectively. Despite experiencing delayed milk spot appearance, slower weight gain following separation, and receiving suboptimal maternal care, ZFP57m-/+ pups exhibited enhanced weight gain during lactation (Figure 4H). Consequently, we decided to investigate the composition of milk from ZFP57^-/-^ and WT dams and performed lipidomic analysis of milk collected on lactation day 8. We found that ZFP57^-/-^ milk contained higher levels oxidised triglycerides (36:0), and lower levels of phosphatidylcholines (38:6 and 34:1) (Figure 4I). Fewer phosphatidylcholines might impact the size of phospholipid-coated lipid droplets in the milk, and potentially body fat accumulation. Previous studies have indicated that dietary phospholipids provided to young mice can alter body fat accumulation^31^. Overall, this indicates that ZFP57^-/-^ mothers produce abnormal milk and that this is associated with growth disparities observed in their pups during lactation, ultimately resulting in enhanced weight gain.

### ZFP57^m-/+^ offspring born to ZFP57^-/-^ dam present with poor life-long health status

Excessive childhood weight gain is recognised as a significant risk factor for cardiovascular and metabolic diseases^32^. To explore the long-term effects of abnormal milk consumption and enhanced growth during lactation, we weighed ZFP57^m-/+^ offspring born to ZFP57^-/-^ dams which consumed abnormal milk and exhibited enhanced growth during lactation, alongside ZFP57^+/-p^ controls from WT dams for up to 6 months. We found that adult ZFP57^m-/+^ born to ZFP57^-/-^ dams continued to accumulate more weight than ZFP57^+/-p^ from WT dams, despite being fed the same chow diet and co-housed (Figure 5A-B). Notably, the weight gain was exacerbated over time (Supplementary Figure 6A). At 6 months, both male and female ZFP57^m-/+^ offspring born to ZFP57^-/-^ dams displayed higher fat mass, lower lean mass and a decreased lean/fat ratio compared to ZFP57^+/-p^ offspring born to WT dams (Figure 5C-E). ZFP57^m-/+^ offspring born to ZFP57^-/-^ dams exhibited impaired glucose tolerance, as indicated by a slower decrease in blood glucose levels during an intraperitoneal glucose tolerance test (IP-GTT) compared to ZFP57^+/-p^ offspring born to WT dams (Figure 5F). However, they performed similarly in an intraperitoneal insulin tolerance test (IP-ITT) (Supplementary Figure 6B-C). We further evaluated the mice using a multi-parameter metabolic assessment system (see Methods). Remarkably, ZFP57^m-/+^ offspring born to ZFP57^-/-^ dams displayed a significantly lower respiratory quotient (Figure 5G), indicating a higher reliance on fat as an energy source compared to ZFP57^+/-p^ offspring born to WT dams. Additionally, the total energy expenditure was lower in ZFP57^m-/+^ compared to ZFP57^+/-p^ offspring (Figure 5H), despite similar levels of ambulatory activity, food consumption and water intake (Supplementary Figure 6C-E). This was supported by higher fat oxidation (Figure 5I) and lower carbohydrate oxidation (Figure 5J) in ZFP57^m-/+^ offspring. Together, our findings indicate that early-life exposure induces metabolic reprogramming in ZFP57^m-/+^ offspring, leading to long-lasting alterations in weight gain, body composition, metabolic rate, and fat oxidation.

**Figure 5.**
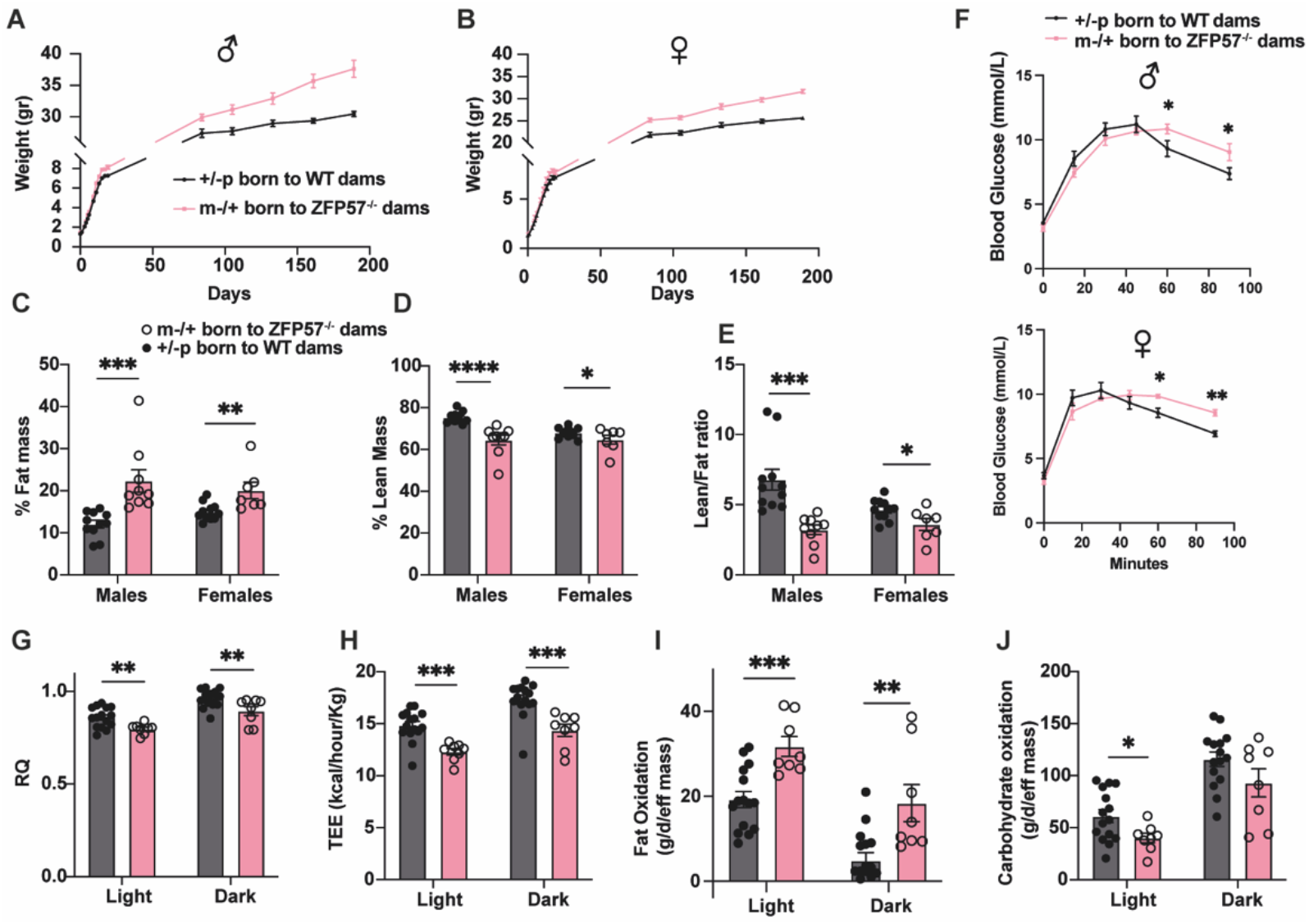
Pups born to ZFP57^-/-^ dams show metabolic syndrome hallmarks during lactation and adulthood. Long-term growth trajectories of male **(A)** and female **(B)** offspring born to ZFP57^-/-^ or WT dams post-weaning, *n* = 46-53 pups from 11-12 litters. **(C-E)** Body composition assessment of male and female offspring of ZFP57^-/-^ and WT dams determined by TD-NMR and normalised to total body weight. **(C)** % Fat mass **(D)** Lean mass, and **(E)** Lean/Fat ratio. *n* = 8-15 pups/ maternal genotype from 2-3 litters. Values are mean ±SEM Two-tailed Student’s *t*-test. **(F)** Glucose tolerance test of offspring born to ZFP57^-/-^ or WT dams at 6 months of age. *n* = 8-15 pups/ maternal genotype from 2-3 litters. Data are mean ±SEM. **(G-J)** Metabolic parameters obtained by monitoring offspring born to ZFP57^-/-^ or WT dams at 6 months of age, by the Promethion high-definition behavioural phenotyping system over 48h period, including **(G)** Respiratory quotient **(H)** Total energy expenditure **(I)** Fat oxidation **(J)** Carbohydrate oxidation. All parameters were normalised to effective mass. Data are mean ±SEM from *n* = 8-15 pups/ maternal genotype from 2-3 litters. Two-tailed Student’s *t*-test. * p<0.05, ** p<0.01, *** p<0.001.

### Cross-fostering ZFP57^m-/+^ and ZFP57^+/-p^ offspring shows synergistic mother-offspring effects during lactation

To attempt to uncouple the maternal-offspring in utero experience from the postnatal growth phenotypes and to further elucidate the effects of maternal genotype on postnatal phenotype, we conducted cross-fostering experiments. WT, ZFP57^m-/+^ and ZFP57^+/-p^ offspring were cross-fostered to either WT or ZFP57^-/-^ dams on postnatal day 1 (Figure 6A). We assessed their weight gain during lactation, using WT pups raised by their WT mothers as a reference. Both WT and ZFP57^+/-p^ offspring showed similar weights at weaning and growth trajectories when raised by or cross-fostered to a WT dam indicating that paternal ZFP57 genotype did not influence postnatal growth trajectory and that being heterozygous *per se*, had no influence on outcome. Consistently with our earlier experiments, ZFP57^m-/+^ pups cross-fostered to ZFP57^-/-^, exhibited substantial weight gain during lactation, however, this weight gain was exacerbated by cross-fostering to a WT dam suggesting that offspring exposed to a ZFP57 mutant in utero environment are more severely compromised if nursed by a mother of a different ZFP57 genotype. Surprisingly, cross-fostering ZFP57^+/-p^ pups to ZFP57^-/-^ dams resulted in a severe failure to thrive, again indicating that pups born to a non-mutant ZFP57 mother are more severely compromised when nursed by a ZFP57 genotypically different mother (Figure 6B, Supplementary Figure 7B, and 8B-C).

**Figure 6.**
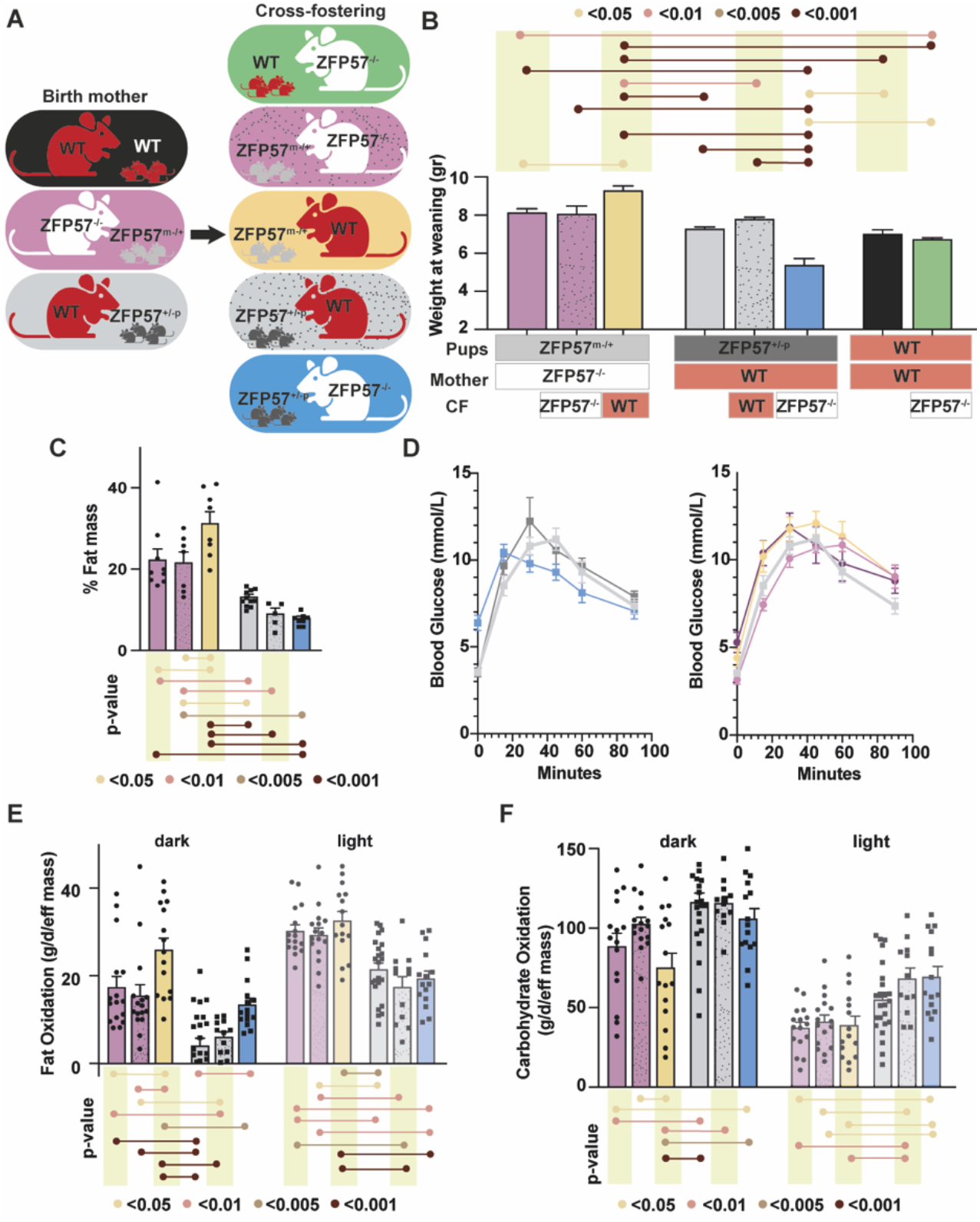
Cross-fostering ZFP57^m-/+^ and ZFP57^+/-p^ offspring show synergistic effects of mother and offspring during lactation with long-lasting implications for energy metabolism. **(A)** Schematic representation of maternal and paternal genotypes of the examined animals, and the genotypes of the foster dams used. **(B)** Male pup weights at weaning in the groups described in A (*n=*24-32/group). Data are mean ± SEM. One-way ANOVA with Tukey’s corrections. **(C)** Fat mass percentage of male ZFP57^+/-p^ and ZFP57^m-/+^ offspring at adulthood. *n*=8-15 pups/group from 2-3 litters. Data are mean ± SEM. One-way ANOVA with Tukey’s corrections. **(D)** Intraperitoneal glucose tolerance test (IP-GTT) of male ZFP57^+/-p^ offspring (left) and ZFP57^m-/+^ offspring (right). ZFP57^+/-p^ offspring born to WT dams is plotted twice for comparison (light grey). *n*=8-15 pups/group from 2-3 litters. Data are mean ± SEM. **(E-F)** Metabolic parameters obtained by monitoring male offspring born to ZFP57^-/-^ or WT dams at 6 months of age, by the Promethion high-definition behavioural phenotyping system over 48h period, including **(E)** Fat oxidation **(F)** Carbohydrate oxidation. All parameters were normalised to effective mass. *n* = 8-15 pups/ group from 2-3 litters. Data are mean ±SEM. One-way ANOVA with Tukey’s corrections. * p<0.05, ** p<0.01, *** p<0.001. **** p<0.001

These experiments highlight the importance of concordance between the genotype of the gestational and nursing mother and emphasises the role of in utero adaptation to postnatal resources provided by the birth mother. Notably, the failure of ZFP57^+/-p^ pups to thrive was observed only when they were cross-fostered to ZFP57^-/-^ dams that differed in genotype from their birth mothers. Additionally, ZFP57^m-/+^ pups which showed the tendency to gain excessive weight when raised by ZFP57^-/-^ dams, exhibited greater weight gain when raised by a WT dam, which differed in genotype from their birth mothers.

### Cross-fostering pups with monoallelic deletion of ZFP57 to a different maternal genotype exacerbates metabolic hallmarks

To investigate the enduring nature or reversibility of the growth phenotypes during lactation, we retained the weaned cross-fostered offspring for 6 months. Subsequently, comprehensive metabolic measurements were conducted. We found that ZFP57^m-/+^ offspring cross-fostered to another ZFP57^-/-^dam show similar body composition as ZFP57^m-/+^ offspring born to ZFP57^-/-^, characterised by high percentage of fat mass, low percentage of lean mass and a reduced lean/fat ratio (Figure 6C, Supplementary Figures 7C-E and 8D-F).

However, the ‘maladapted’ ZFP57^m-/+^ offspring cross-fostered to WT dams which showed increased weight gain during lactation, displayed an exacerbated obesity phenotype at 6 months of age, with a significantly higher percentage of fat mass when compared with ZFP57^m-/+^ raised by ZFP57^-/-^ dams (Figure 6C, Supplementary Figure 8D). In contrast, the ‘maladapted’ ZFP57^+/-p^ offspring, which failed to thrive when cross-fostered to ZFP57^-/-^ dams were able to catch up post-weaning, without long-term effects on body composition (Supplementary Figures 7C-D, 8D-F).

Male, but not female ZFP57^+/-p^ offspring cross-fostered to ZFP57^-/-^ dams showed significantly higher fasting glucose levels compared to ZFP57^+/-p^ offspring raised by WT dams (Supplementary Figures 7E, 8G). When challenged with IP-GTT, ‘maladapted’ ZFP57^m-/+^ offspring cross-fostered to WT dams performed worse than ZFP57^m-/+^ offspring raised by ZFP57^-/-^ dams. These cross-fostered offspring took longer to normalise the levels of the injected glucose, supported by a larger value for the area under the curve (AUC) (Figure 6D, Supplementary Figures 7F, 8H-I). IP-GTT revealed intriguing dynamics in the ‘maladapted’ ZFP57^+/-p^ offspring cross-fostered to ZFP57^-/-^ dams which failed to thrive as pups. These animals showed peak glucose levels 15 minutes post-injection, followed by a sharp decrease. In contrast, ZFP57^+/-p^ offspring raised by WT dams continually increased glucose levels up until 30- or 45-minutes post-injection, followed by clearance of the glucose and normalisation of the levels (Figure 6D).

Further metabolic status profiling of the cross-fostered offspring highlighted that ZFP57^m-/+^ offspring cross-fostered to WT dams exhibited distinct metabolic differences from the same offspring raised by or cross-fostered to ZFP57^-/-^ dams. During the dark period, ZFP57^m-/+^ offspring cross-fostered to WT dams (hence differing in genotype from their birth mothers) showed significantly higher fat oxidation compared to offspring with the same epigenotype raised by or cross-fostered to ZFP57^-/-^ dams (Figure 6E), while their carbohydrate oxidation was much lower in the dark (Figure 6F). Additionally, during the dark period, ZFP57^m-/+^ offspring cross-fostered to WT dams showed significantly lower respiratory quotient, (Supplementary Figure 7G), indicating higher utilisation of fat as an energy source compared to ZFP57^m-/+^ offspring raised by or cross-fostered to ZFP57^-/-^ dams. The total energy expenditure of these animals was higher than ZFP57^m-/+^ offspring raised by or cross-fostered to ZFP57^-/-^ dams (Supplementary Figure 6H). These findings indicate that ZFP57^m-/+^ offspring cross-fostered to WT dams whose genotype was different from their birth mother, showed an inferior metabolic profile than those raised by ZFP57^-/-^ dams whose genotype was, or represented, that of their birth mother.

Examination of the metabolic performance of ZFP57^+/-p^ offspring cross-fostered to ZFP57^-/-^ dams revealed that their respiratory quotient, total energy expenditure and carbohydrate oxidation were comparable to those of ZFP57^+/-p^ offspring raised by or cross-fostered to WT dams (Figure 6F, Supplementary Figure 7G-H). However, they did exhibit increased fat oxidation during the dark period (Figure 6E), suggesting that despite experiencing significant developmental challenges as pups nurtured by a mother genotypically different from their birth mother, the metabolic parameters of these animals largely normalise post weaning.

## Discussion

In our study, we have identified ZFP57 as a key modulator of postnatal nutritional resources, specifically affecting mammary gland development and milk production.

Previous studies have suggested that imprinting may be crucial in the mammary gland^2^, with some demonstrating functional roles for imprinted genes in this tissue^3,33^. However, although ZFP57 is a major regulator of imprinting, we show that it does not act through imprints in the mammary gland. This is supported by the absence of reciprocal changes in imprinted gene expression associated with loss of imprinting e.g. increased H19 and decreased Igf2. Instead of a complete loss of one gene and a doubling of expression in the other, both genes are upregulated. Moreover, only a small number of imprinted genes display differential expression. Additionally, most of the DEGs were observed in luminal progenitor cells, which express very low levels of ZFP57. Overall, this suggests that ZFP57 is not acting directly in the mammary gland, but influences the tissue indirectly. This could be attributed to the developmental effects of ZFP57, a transcriptional defect, inter-organ communication or secondary effects of other genes.

Analysis of the enriched pathways among the DEGs in mammary tissue highlighted that the top enriched terms were related to mammary gland development, milk composition and body weight, features that are affected in ZFP57^-/-^ dams or their offspring. This suggests that ZFP57 controls a specific set of mammary- and milk-related genes consequently influencing these processes.

Our model suggests that ZFP57 regulates the provision of nutritional resources to offspring through two distinct mechanisms – prenatally, through genomic imprinting and postnatally, through mammary gland and lactation-related genes.

A limitation of our study is the absence of Cre-line specifically deleting ZFP57 in mammary cells. However, such a model will prevent us from examining the systemic physiological effects of ZFP57, since other vital organs that communicate with the mammary gland during pregnancy and lactation, such as the brain, the pituitary gland, the ovaries and placenta will retain their ZFP57 levels.

Our study reveals a novel finding challenging the prevailing notion of environmental co-adaptation between mother and offspring. Instead, we demonstrate that this co-adaptation is genetically driven. Through our cross-fostering experiments, we observed that genetically identical offspring, raised by dams with a different genotype to their birth mother, exhibit more extreme phenotypes. Specifically, ZFP57^+/-p^ fail to thrive during lactation when they are raised by ZFP57^-/-^ dams which produce abnormal milk, while ZFP57^m-/+^ raised by WT dams show exacerbated metabolic syndrome hallmarks and obesity. Given that these crosses exclusively result in heterozygous offspring, it is impossible to exclude the possibility that the genotype of the pups might play a role in their mal-adaptation to a foster mother of a different genotype. These findings suggest that these offspring experience maladaptation due to maternal genetic factors rather than environmental influences.

The precise factors contributing to this genetic adaptation remain an open question. Exploring factors such as altered body composition in ZFP57^m-/+^ and the failure to thrive in ZFP57^+/-p^, potentially related to gene dosage and haploinsufficiency in the father’s sperm, would be intriguing avenues for further investigation. Additionally, understanding the relative contributions of in-utero and ex-utero exposure, as well as the specific impacts of milk composition, requires further work.

Overall, this study enhances our understanding of mother-offspring co-adaptation and its long-term effects on the lifecourse. It sheds new light on the interplay between genetic factors and early-life nutrition, expanding our knowledge of this complex relationship.

## Author contributions

ACFS and GH planned the experiments and interpreted the data. GH performed all the expriments, analysed the data, wrote the manuscript and prepared the figures. All authors contributed to the writing and editing of the manuscript. BA performed Tunel stainings, immunohistochemistry, confocal microscopy, image analysis, quantitative PCR and assisted with metabolic assays. KRC performed RNA-seq analysis. HT performed RNA-seq differential expression analysis. NT collected and analysed embryonic imprinted gene expression. LAM analysed expression in existing datasets. AKS performed quantitative PCR, contributed to suckling assays and assisted with metabolic assays. SP assisted with RNA processing and library preparations. BJ and AK analysed milk composition.

## Acknowledgements

This work was funded by MRC grants MR/R009791/1 to ACF-S and MR/W003783/1 to ACF-S and GH. G Hanin was supported by the Royal Society Newton Fellowship and FEBS long-term fellowship. B Alsulaiti was supported by Qatar National Research Fund (GSRA8-I-1-0505-21030).

The Cambridge NIHR BRC Cell Phenotyping Hub, and the Cambridge Advanced Imaging Centre contributed support to the project. We thank Dr Leila Muresan for assisting in image quantification and analysis.

