## Supplementary data for "ZFP57 is a regulator of postnatal growth and life-long health"

### Methods

##### Animals

Mouse work and the experiments for this study were approved by the University of Cambridge Animal Welfare, and Ethical Review Body and performed under the UK Home Office Animals (Scientific Procedures) Act 1986. (Home Office project licence # PC213320E and PP8193772).
ZFP57 mutants[1] were backcrossed to 129aa background for >30 generations and housed under a 12h light / 12h dark photocycle 22°C air temperature and 21% oxygen saturation with access to water and standard laboratory chow diet ad libitum (RM3, Special Dietary Services [SDS], Witham, UK) (11% fat, 7% simple sugars of energy contribution [%kcal]).
Mice were mated at 12-16 weeks of age, embryos were collected at E12.5, and adult tissues including the hypothalamus, pituitary glands and mammary glands were collected from 12-14 week-old females. Nulliparous mice were oestrus cycles matched.

##### *Cross fostering*

Heterozygous pups born to either ZFP57$^{-/-}$ x WT (mat x pat) or WT x ZFP57$^{-/-}$ (mat x pat) crosses within 24 hours of each other were used for cross-fostering. The original litter and the dam were removed from the dam's cage, and the cross-fostered pups were gently rolled in the nesting material of the fostering dam. Subsequently, the original dam was returned to her home cage with a new litter, which was weighed every 1-2 days until weaning.
The number of pups was normalised to the size of the smaller litter.

##### *Milk let-down*

Pups and dams were separated at postnatal day 9 for 4 hours and were placed in proximity to each other in a heated cabinet. Following separation, the dam and pups were weighed using analytical scales immediately before their return to the nest (time 0), and at intervals of 30 minutes up to 3 hours thereafter.

##### *Maternal behaviour*

Dam behaviour was quantified in their home cage based on several parameters, including the quality of the built nest, scored 0-5, time to approach pups following their weighing, and time to retrieve pups and place them back in the nest[2]

##### *Mouse Milking*

Mouse milking was performed on lactation day 8, on dams with at least 4 pups/litter. Pups and dams were separated for 4 hours and placed in proximity to each other in a heated cabinet. Following the separation, dams were weighed and injected intraperitoneally with Ketamine (Ketavet, Zoetis, UK) 90 mg/Kg and Xylazine (Xylacare, Animalcare, UK) 10 mg/Kg in Saline, and 2 IU/Kg Oxytocine (sigma O4375-1000IU) in DDW. Milk was expressed manually using a gentle massage and collected using a P20 pipette.

##### *Multi-parameter metabolic assessment*

TD-NMR (LF50H Minispec, Bruker, Coventry UK) was used to determine longitudinal changes in body composition (lean and fat mass). Metabolic and activity profiles of the mice were assessed using the Promethion High-Definition Behavioral Phenotyping System (Sable Systems, Las Vegas, NV, USA) as described previously[3]. Data analysis was performed using ExpeDATA (Sable Systems, Las Vegas, NV, USA). The Respiratory quotient (RQ) was calculated as the ratio of VCO2/VO2. Total energy expenditure (TEE) was calculated as VO2 x (3.815 + 1.232 x RQ), normalised to effective body mass calculated by ANCOVA, and expressed as kcal/h/kg eff. Mass. Fat oxidation (FO) was calculated as *FO* = 1.69 x *VO2* – 1.69 x *VCO2* and expressed as g/d/kg eff. Mass. Ambulatory activity and position were monitored simultaneously with the collection of the calorimetry data using XYZ beam arrays with a beam spacing of 0.25 cm.

##### *Glucose tolerance tests (GTT)*

For intraperitoneal GTT (IP-GTT), mice fasted for 16 hours and were then injected with 10% glucose (D-glucose, Sigma Aldrich, MO, USA) at a 2 mg/kg dose in double distilled water. Mice were bled from a tail clip. Blood glucose was measured before injection (time 0) and 15, 30, 60, 90 and 120 minutes post-injection using a handheld glucometer (Accu-Chek Performa, Roche. Mannheim, Germany). Blood glucose levels were compared between Heterozygous offspring born to either ZFP57-/- x WT (mat x pat) or WT x ZFP57-/- (mat x pat) crosses, at each time point using multiple unpaired t-tests corrected for multiple comparisons.

##### *Genotyping*

ZFP57$^{-/-}$ mice were genotyped using the following primers: forward knockout, GAAAGTCCTGAATGCGTTGC, reverse knockout, GTGGGAAAGGGTTCGAAGTT, and ZFP57 forward wildtype AGGACGTGGCAGTGTCTTTC and ZFP57 reverse wildtype GACAAATGTCAGGTTCTTGAA. PCR product levels were then determined using agarose gel electrophoresis.

##### ELISA

Circulating progesterone and oestradiol were determined in serum, using 17-beta-Oestradiol ELISA (Abcam ab108667), and progesterone ELISA (Enzo Life Sciences, ADI-900-001) according to manufacturer’s instructions.

##### Flow cytometry

Single-cell suspensions of mammary cells were prepared from WT or ZFP57$^{-/-}$ females at three different stages: nulliparous, gestation day 9.5 or lactation day 2. Pregnancy was confirmed during dissections. Mammary fragments were digested for 1 hour at 37°C using Krebs–Ringers – HEPES, 2.5 mM glucose (Sigma-Aldrich, D9434), 2% fetal bovine serum (Gibco, 16-000-044), 200 µM adenosine (Sigma-Aldrich, A9251), 1 mg/ml collagenase (Sigma-Aldrich, C2139), pH=7.4. Single cells were then filtered, and blocked with 10% normal rat serum (Sigma-Aldrich R9759-10ML) for 15 minutes at 4°C and stained with the following antibodies: Anti-Mouse CD45 APC-eFluor780 (Invitrogen, 47-0451-82), Anti-Mouse CD31 PE-Cy7 (Invitrogen, 25-0311-82), Anti-Mouse TER-119 Biotin (Invitrogen, 13-5921-82), Anti-Mouse BP-1 Biotin (Invitrogen, 13-5891-81), Anti-Mouse EpCam Alexa Fluor647 (Biolegend, 118211), Anti-Mouse CD49b PE (Biolegend, 103506), Anti-Mouse CD49f Alexa Fluor488 (Biolegend, 313608). Viability was assessed using Zombie Aqua dye (Biolegend 423101).

Subsequently, cell populations were sorted using BD ARIA II cell sorter (BD Biosciences). Data were analysed using FlowJo 10.8.1.

**RNA extraction**

RNA was extracted from sorted cells or whole mammary gland tissue using miRNeasy micro kit (QIAGEN, 217084) or TRI reagent (Sigma-Aldrich, St. Louis, MO, USA) respectively, according to standard protocols. RNA concentration was determined using Qubit Fluorometer (ThermoFisher), and RNA integrity was quantified using the 2100 Bioanalyzer instrument (Agilent).

**mRNA quantifications**

mRNA levels were determined by quantitative reverse transcription PCR (RT-qPCR) using the RevertAid H Minus First Strand cDNA Synthesis kit (Thermo Scientific) followed by quantification in technical triplicates with Brilliant III Ultra-Fast SYBR® Green QPCR Master Mix (Agilent) on the LightCycler 480 Instrument (Roche). Relative expression was normalised to β-Tubulin expression unless otherwise noted and calculated using the ΔCt method.
For ChIP-qPCR, no template controls and 10% ChIP input were run alongside the ZFP57 and Rabbit IgG ChIP samples for each primer pair. Enrichment was calculated as per cent input. Primers were designed using Primer3 software. Primer sequences are listed in Supplementary Table 1

**Library generation and RNA sequencing**

Libraries for RNA sequencing (RNA-seq) from sorted mammary cells were generated using SMARTer stranded total RNA-seq kit V2 – Pico input mammalian (TaKaRa bio, 634414) according to the manufacturer's instructions. The quality and RNA integrity number were assessed using Bioanalyzer 2100 (Agilent), Qubit fluorometer (Thermo Fisher) and sequenced using the NovaSeq 6000 system (Illumina).

RNA-seq processing pipeline was built using Snakemake v6.10.0[4]. Reads were trimmed using Cutadapt v3.4[5] with the following settings: Illumina TruSeq adapters, -U4 --quality-cutoff 20,

minimum length 50 base pairs. Read quality was assessed before and after trimming with FastQC v0.11.9 (Andrews 2010, http://www.bioinformatics.babraham.ac.uk/projects/fastqc). Further quality control metrics were obtained by aligning the reads to the standard GRCm38/mm10 mouse reference genome (Ensembl release 102) using STAR v2.7.9a[6], followed by RSeQC v4.0.0[7] for quality alignment control. Quality metrics were compiled using MultiQC v1.10.1[8]. Transcript quantification was performed using Salmon v1.5.0 in mapping-based mode[9]. Read count matrices were prepared using the R/Bioconductor package tximport v1.20.0[10]. DESeq2 v1.32.0[11] was used to detect differentially expressed protein-coding genes with at least 20 reads, false discovery rate (FDR) threshold of 0.05 and an absolute log2 fold-change threshold of 1.0. Differential expression gene analysis was performed separately for each cell type and stage. The DESeq2 package was also used to obtain normalised read counts using the variance stabilising transformation method.
UMAPS[12] were generated using normalized read counts obtained from DEseq2's "variance stabilising transformation" method. UMAPs were made using 9 neighbours, with a min distance of 0.25, and using the Euclidean metric. Gene ontologies of differentially expressed genes were completed using Enrichr[13] , and the top 4 most enriched terms were selected for visualization.

**Milk lipidomic analysis**

All solvents and additives were of HPLC grade or higher and purchased from Sigma Aldrich (Haverhill, Suffolk, UK) unless otherwise stated.

The protein-precipitation liquid extraction protocol has been described previously[14]. Briefly, 10 µL of mouse milk was transferred into a 2 mL screw cap Eppendorf plastic tube (Eppendorf, Stevenage, UK). Immediately, 650 µL of chloroform was added to each sample, followed by thorough mixing. Then, 100 µL of the LIPID-IS (5 µM in methanol), 100 µL of the CARNITINE-IS (5 µM in methanol) and 150 µL of methanol was added to each sample, followed by thorough mixing. Then, 400 µL of acetone was added to each sample. The samples were vortexed and centrifuged for 10 minutes at ~20,000 g to pellet any insoluble material. The supernatant was pipetted into separate 2 mL screw cap amber-glass auto-sampler vials (Agilent Technologies, Cheadle, United Kingdom). The organic extracts were dried down to dryness using a Concentrator Plus system (Eppendorf, Stevenage, UK) run for 60 minutes at 60 degree Celsius. The samples were reconstituted in 100 µL of 2: 1: 1 (propan-2-ol, acetonitrile and water, respectively) then thoroughly vortex. The reconstituted sample was transferred into a 250 µL low-volume vial insert inside a 2 mL amber glass auto-sample vial ready for liquid chromatography with mass spectrometry detection (LC-MS) analysis.
Full chromatographic separation of intact lipids was achieved using Shimadzu HPLC System (Shimadzu UK Limited, Milton Keynes, United Kingdom) with the injection of 10 µL onto a Waters Acquity UPLC® CSH C18 column (Waters, Hertfordshire, United Kingdom); 1.7 µm, I.D. 2.1 mm X 50 mm, maintained at 55 degrees Celsius. Mobile phase A was 6:4, acetonitrile and water with 10 mM ammonium formate. Mobile phase B was 9:1, propan-2-ol and acetonitrile with 10 mM ammonium formate. The flow was maintained at 500 µL per minute through the following gradient: 0.00 minutes_40% mobile phase B; 0.40 minutes_43% mobile phase B; 0.45 minutes_50% mobile phase B; 2.40 minutes_54% mobile

phase B; 2.45 minutes_70% mobile phase B; 7.00 minutes_99% mobile phase B; 8.00 minutes_99% mobile phase B; 8.3 minutes_40% mobile phase B; 10 minutes_40% mobile phase B. The sample injection needle was washed using 9:1, 2-propan-2-ol and acetonitrile. The mass spectrometer used was the Thermo Scientific Exactive Orbitrap with a heated electrospray ionization source (Thermo Fisher Scientific, Hemel Hempstead, UK). The mass spectrometer was calibrated immediately before sample analysis using positive and negative ionization calibration solution (recommended by Thermo Scientific). Additionally, the mass spectrometer scan rate was set at 4 Hz, giving a resolution of 25,000 (at 200 m/z) with a full-scan range of m/z 100 to 1,800 with continuous switching between positive and negative mode.
Data processing—The instrument responses of the analytes were normalized to the relevant internal standard response (producing relative concentrations), these relative concentrations are corrected for the intensity for any extraction and instrument variations.

**Immunoblots**

Samples were lysed in a solution containing 10mM Tris HCl pH=7.4, 1 M NaCl, 1 mM EGTA, and 1 % TX-100, and homogenised with a Kontes pellet pestle, incubated on ice for 10 minutes, centrifuged at 17,900 rcf, 4°C, for 30 minutes, and the clear lysate collected. 40μg protein samples were separated by standard SDS-PAGE procedures on 4-20% Mini-PROTEAN TGX gel (BioRad; 4561095) and ran at 90-130V. After transfer to PVDF membrane (BioRad, 1704156EDU) using the BioRad TransBlot Turbo system (BioRad, 1704150ED)U, and blocking in 5% skimmed milk overnight at 4°C, proteins were visualised using primary antibodies against ZFP57 (Abcam; ab45341; 1:250) and Vinculin (Abcam; ab129002; 1:5000) for 48 hours at 4°C. The membrane was washed with TBST for 30 mins, followed by HRP-conjugated Goat anti-Rabbit secondary antibody incubation (Agilent Technologies; P044801-2; 1:10,000) for 2 hrs at RT, followed by a 30 min wash with TBST. Membrane was developed with SuperSignal™ West Femto Maximum Sensitivity Substrate kit (ThermoFisher; 34095) and exposed for 5 minutes on a Licor imager. Bands were quantified and analysed using densitometry with Fiji software.

**Immunohistochemistry**

Paraffin slides were rehydrated by washing in xylene and decreasing ethanol concentrations in water. Heat-induced antigen retrieval was done by boiling slides for 10 minutes in 10mM citrate buffer pH 6. After washing, slides were incubated with 150 µl/slide of blocking buffer (4% horse serum, 0.05% TWEEN20 and 0.3% Triton X-100) for 60 min, followed by overnight incubation at 4°C with primary antibody diluted in the blocking buffer. Slides were then washed and incubated with fluorescently- conjugated secondary antibodies for 2 h. Nuclear staining was done using 40,6-diamidino-2-phenylindole (DAPI, 5μg/μl for 5 minutes).
Primary antibodies used were anti-K8 (DSHB; TROMA-I; 1:50), anti-αSMA (Novus Bio; NB300-978; 1:200), and anti-Ki67 (Cell Signaling; 12202; 1:200). Stained tissues were imaged using Leica SP8 confocal microscope.
Quantification was performed using Fiji software, using the free-drawing tool, ducts were circled and the intensity of Ki67 was measured, then normalised to DAPI.

**TUNEL staining**

Mammary glands were fixed and sectioned at Nulliparous, gestation days 4.5, 9.5 and lactation day 2, and analysed for apoptosis using terminal deoxynucleotidyl transferase Br-dUTP nick end labelling (Abcam; ab66110) according to the manufacturer's instruction and using a 1:40 secondary antibody dilution. Z-stack images were taken on a Leica SP8 confocal microscope and analysed using FiJi software using the StarDist plugin to identify mammary ducts, calculating the percentage of BrdU intensity within the duct. Analysis included 3-5 animals per stage and genotype, and 27-60 fields per sample.

**Histology**

Whole mounts of mammary tissues from nulliparous mice, gestation and lactation stages we fixed in 4% Paraformaldehyde at 4°C, rinsed in Phosphate-buffered saline and stained overnight with carmine alum solution (Sigma-Aldrich C-1022, Aluminum Potassium sulfate (Sigma Aldrich A-7167). Whole mounts were dehydrated and mounted for imaging using Zeiss Imager Apotome. Morphometric analysis was analysed using Fiji software.

**Statistical analysis**

Flow cytometry, milk lipidomic analysis and pup retrieval assay were performed blinded to genotype. All statistical analyses were performed using GraphPad Prism 8 Software. Statistical significance between two groups was determined by Mann-Whitney tests or two-tailed unpaired t-tests with Welch correction. Statistical significance between multiple groups was performed using one- or two-way ANOVA followed by Tukey's multiple comparisons tests, as appropriate.
Differences were considered significant when *$P < 0.05$, **$P < 0.01$, ***$P < 0.001$, ****$P < 0.0001$. Data are presented as mean ± SEM.
The numbers of samples or litter used for each experiment are indicated in figure legends.

#### Supplementary figures

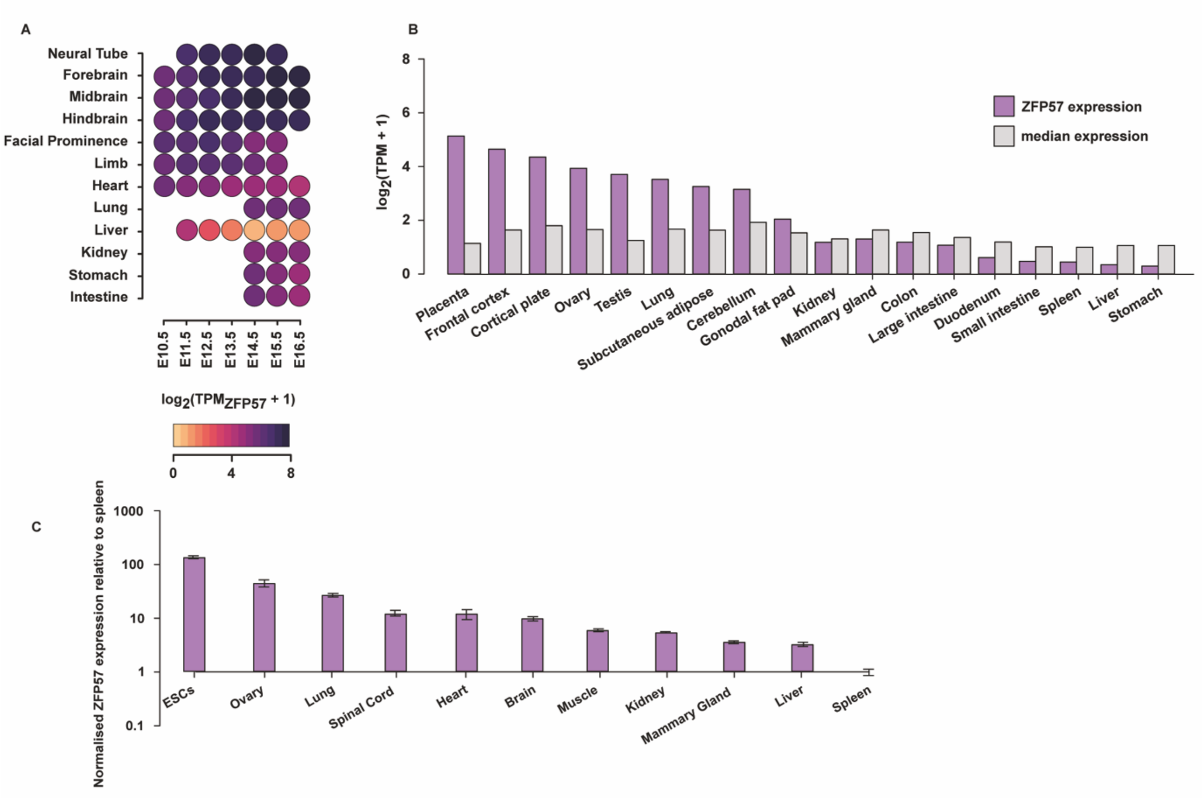


**Supplementary Figure 1 – *ZFP57* expression across tissues and stages**

**(A)** *Zfp57* expression during embryonic development across different tissues and stages based on ENCODE PolyA RNA-Seq data. Expression is measured using TPM, and transformed onto a log2 scale. Darker colour represents higher expression. **(B)** Zfp57 expression in 8-week-old mice across different tissues based on ENCODE RNA-Seq data. Purple bars indicate the expression of Zfp57 measured by TPM and log2 transformed for plotting. Grey bars represent median expression for comparison. **(C)** Relative ZFP57 expression determined by qPCR in adult mouse tissues from C57BL6/J mice. Values are normalised to β-Tubulin. *n*=4-5.

**Supplementary Figure 2 – Flow cytometry gating strategy**

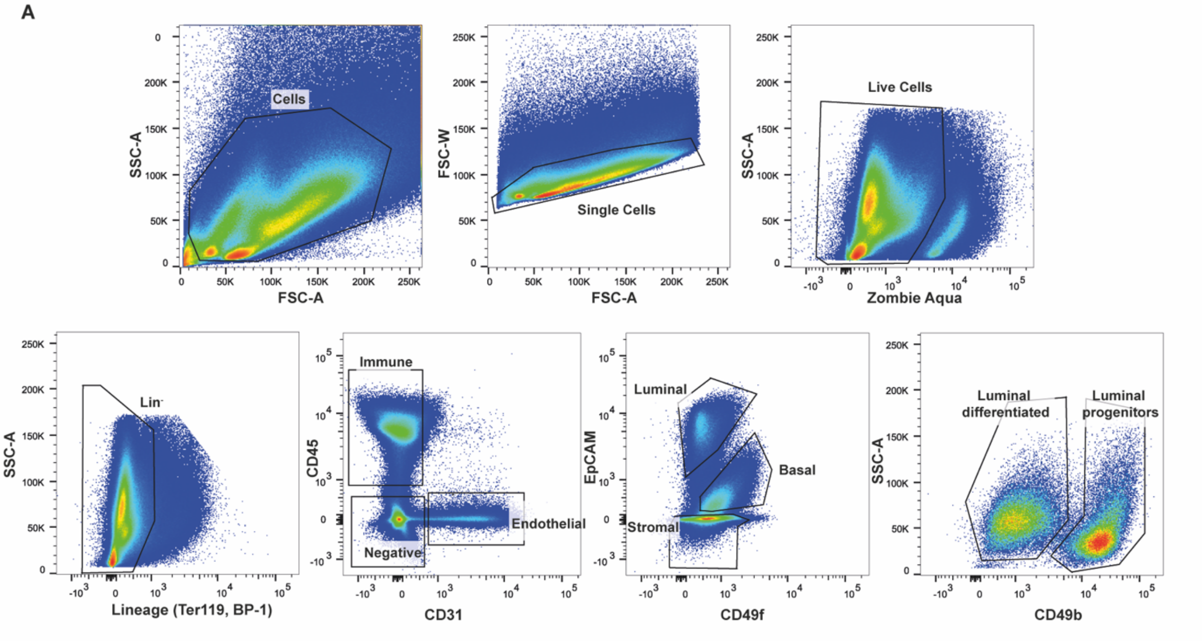


**(A)** Representative flow cytometry plots showing the gating strategy to sort various mammary cell populations. Mammary cells were sorted to detect immune cells (CD45$^{+}$), endothelial cells (CD31$^{+}$), basal cells (LinCD31$^{neg}$CD45$^{neg}$EpCAM$^{lo}$ CD49f$^{hi}$), differentiated luminal cells (Lin$^{-}$CD31$^{neg}$CD45$^{neg}$EpCAM$^{high}$ CD49f$^{low}$ CD49b$^{low}$), and luminal progenitor cells (Lin$^{-}$CD31$^{neg}$CD45$^{neg}$EpCAM$^{high}$ CD49f$^{low}$ CD49b$^{high}$).

**Supplementary Figure 3 – Flow cytometry and abnormal numbers of immune cells in ZFP57$^{-/-}$ mice.**

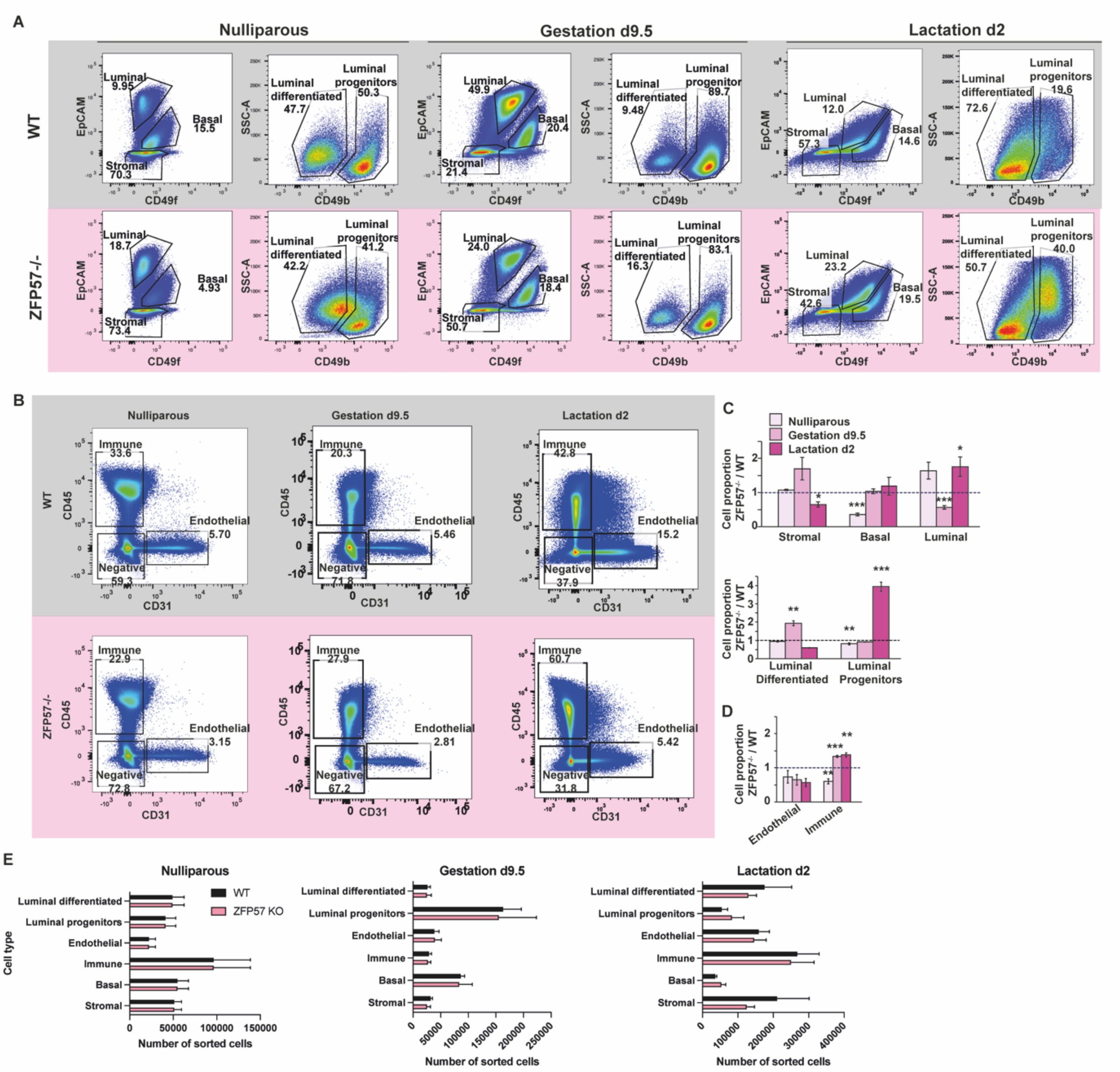


**(A-B)** Representative flow cytometry plots comparing the relative percentages of stromal, basal and luminal (progenitors and differentiated cells) **(A)**, as well as immune and endothelial cell **(B)**. Horizontal panels represent WT (grey) and ZFP57$^{-/-}$ (pink) mice sorted at nulliparous, gestation day 9.5 and lactation day 2 stages. **(C-D)** Quantification of the cellular proportions for each sorted population. Data are presented as the ratio between ZFP57$^{-/-}$/WT across cell types and stages. *n*=5-8/group, Data are mean ± SEM. One-way ANOVA with Tukey's corrections. **(E)** Absolute numbers of cells sorted from WT and ZFP57$^{-/-}$ across cell types and stages. *n*=5-8/group, Data are mean ± SEM. * p<0.05, ** p<0.01, *** p<0.001. *** p<0.001.

##### Supplementary Figure 4 – RNA-seq

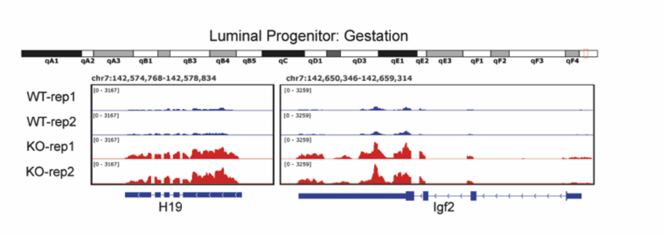


Genome screenshot of RNA-seq from WT and ZFP57$^{-/-}$ mice at the Igf2-H19 locus which are differentially expressed in Luminal progenitor cells during gestation d9.5.

##### Supplementary Figure 5 – Characteristics of ZFP57$^{-/-}$ dams and their maternal behaviours

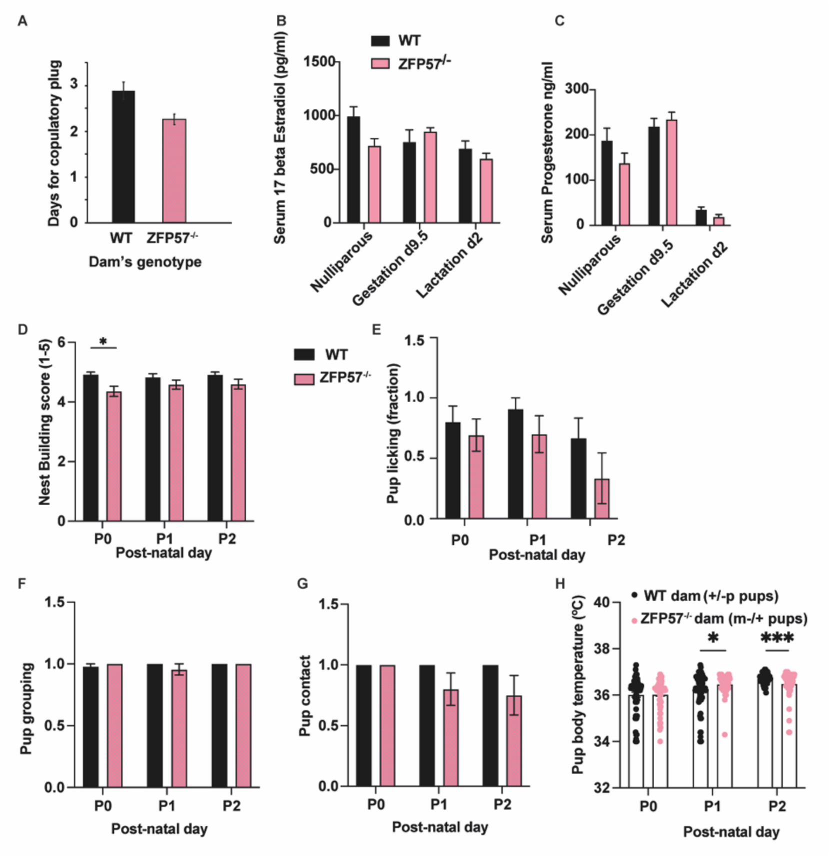


**(A)** Typical duration for a copulatory plug that resulted in pregnancy to be observed. n=117-128/genotype. Values are mean ±SEM. **(B-C)** Circulating hormone levels in WT and ZFP57$^{-/-}$ nulliparous dams and during pregnancy and lactation, *n*=5/genotype and stage. **(B)** Serum 17-β-estradiol **(C)** Serum progesterone. Values are mean ±SEM. **(D-H)** Quantification of maternal-pup interaction phenotypes between P0 and P2: **(D)** Nest building **(E)** Pup licking **(F)** Pup grouping **(G)** Pup contact **(H)** Pup body temperature. *n*=74-93 pups/group from 13-14 litters. Values are mean ±SEM. Two-tailed Student's *t*-test. * p<0.05, ** p<0.01, *** p<0.001. *** p<0.001.

**Supplementary Figure 6 – Additional metabolic parameters of pups born to ZFP57$^{-/-}$ dams during adulthood.**

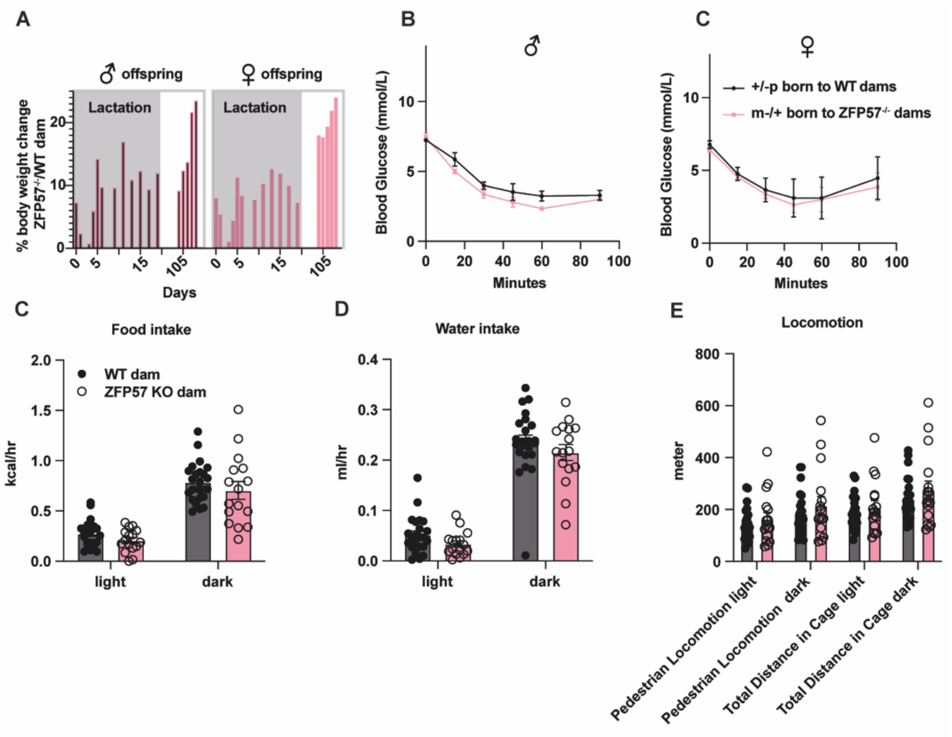


**(A)** Body weight change between offspring of ZFP57$^{-/-}$ and WT dams throughout adulthood, represented as the ratio between the two genotypes. *n*=46-53 pups from 11-12 litters/genotype. Data are delta of mean ± SEM. **(B-C)** Insulin tolerance test of male **(A)** and female **(B)** offspring born to ZFP57$^{-/-}$ or WT dams at 6 months of age. *n*=3-5 pups/group. **(C-E)** Metabolic parameters obtained by monitoring offspring born to ZFP57$^{-/-}$ or WT dams at 6 months of age, by the Promethion high-definition behavioural phenotyping system over 48h period, including **(C)** Food intake **(D)** Water intake **(E)** Locomotion, displayed as pedestrian locomotion and total distance. Data are mean ±SEM from *n* = 8-15 pups/gorup from 2-3 litters. Two-tailed Student's *t*-test. * p<0.05, ** p<0.01, *** p<0.001.

**Supplementary Figure 7 - additional data on cross-fostering male ZFP57$^{m-/+}$ and ZFP57$^{+/-p}$ offspring.**

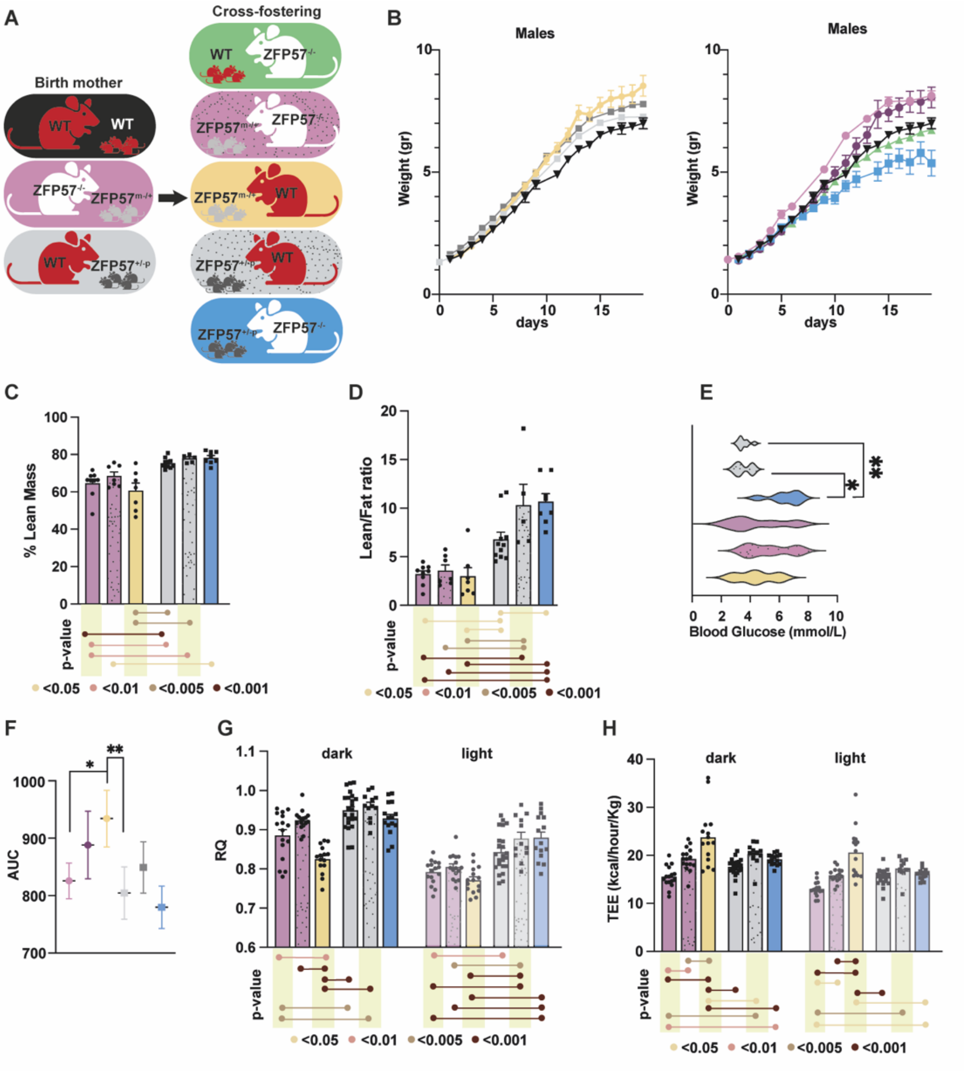

**(A)** Schematic representation of maternal and paternal genotypes of the examined animals, and the genotypes of the foster dams used.
**(B)** Growth trajectories during the lactation period, of male pups raised by WT (left) or ZFP57$^{-/-}$ dams (right)**. Left -** WT pups raised by WT dams (black triangles) served as a baseline, compared to ZFP57$^{+/-p}$ pups raised by WT dams (light grey squares) or cross-fostered to WT dams (dark grey squares), and ZFP57$^{m-/+}$ (yellow circles) pups raised by or cross-fostered to WT dams. **Right -** WT pups raised by WT dams (black triangles) served as a baseline, compared to WT pups raised by ZFP57$^{-/-}$ dams (green triangles), ZFP57$^{m-/+}$ pups raised by ZFP57$^{-/-}$ dams (light purple circles) or cross-fostered to ZFP57$^{-/-}$ dams (dark purple circles), and ZFP57$^{m-/+}$ (blue squares) pups raised by or cross-fostered to ZFP57$^{-/-}$ dams. (*n*=24-32/group). **(C-D)** TD-NMR measurements show body compositions of male ZFP57$^{+/-p}$ and ZFP57$^{m-/+}$ offspring, including, lean mass percentages **(C)**, and lean/fat ratio **(D)**. *n*=8-15 pups/group from 2-3 litters. Data are mean ± SEM. One-way ANOVA with Tukey's corrections. **(E)** Fasting glucose of ZFP57$^{+/-p}$ and ZFP57$^{m-/+}$ male offspring. *n*=8-15 pups/group from 2-3 litters. Data are mean ± SEM. One-way ANOVA with Tukey's corrections. **(F)** Area under the curve quantifying the performance of different groups of

males in the IP-GTT in Figure 6D. *n*=8-15 pups/group from 2-3 litters. Data are mean ± SEM. **(G-H)** Metabolic parameters obtained by monitoring male offspring born to ZFP57$^{-/-}$ or WT dams at 6 months of age, by the Promethion high-definition behavioural phenotyping system over 48h period, including **(G)** Respiratory quotient **(H)** Total energy expenditure. All parameters were normalised to effective mass. Data are mean ±SEM from *n* = 8-15 pups/ maternal genotype from 2-3 litters. * p<0.05, ** p<0.01, *** p<0.001.

**Supplementary Figure 8 - additional data on cross-fostering female ZFP57$^{m-/+}$ and ZFP57$^{+/-p}$ offspring.**

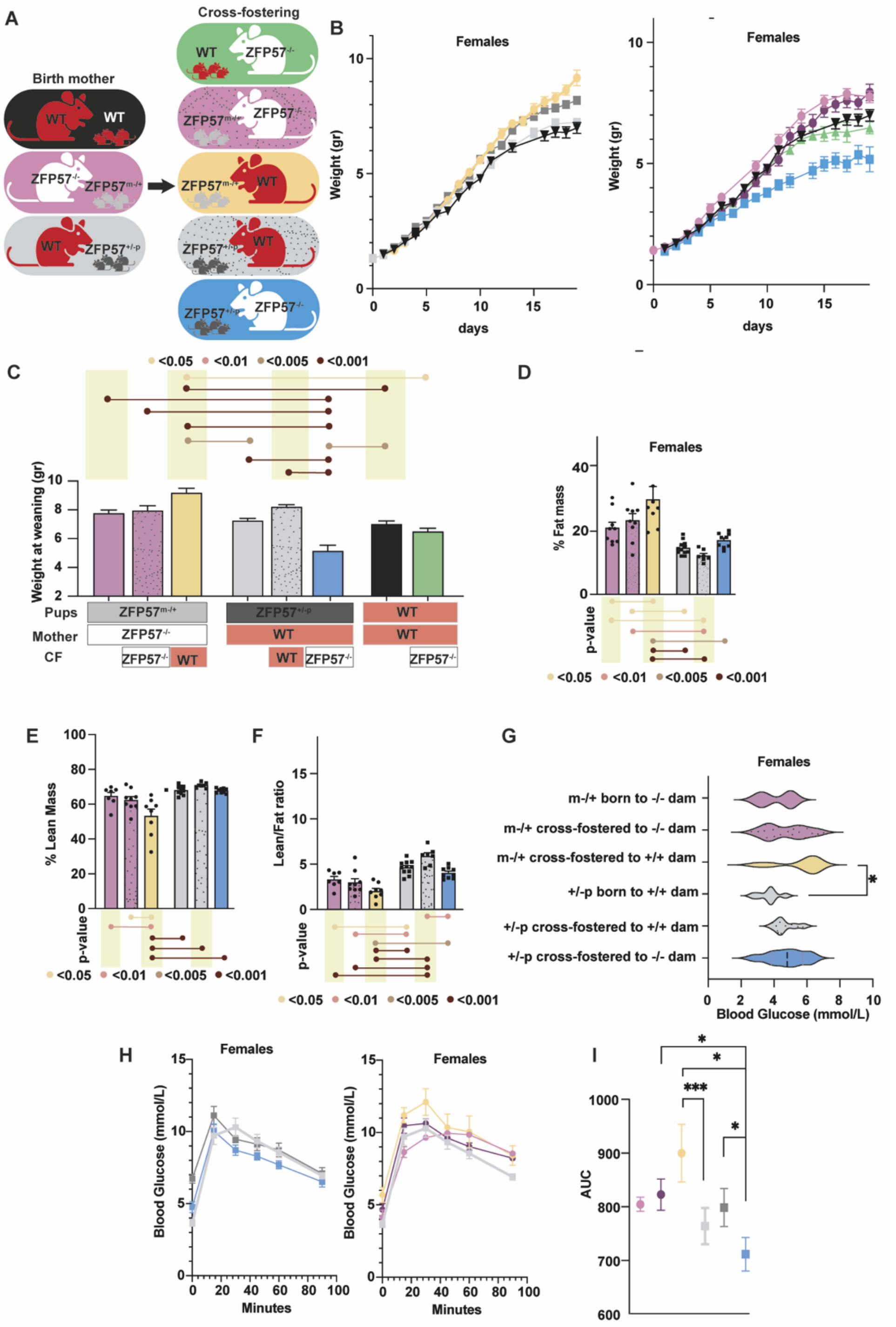


**(A)** Schematic representation of maternal and paternal genotypes of the examined animals, and the genotypes of the foster dams used.

**(B)** Growth trajectories during the lactation period, of female pups raised by WT (left) or ZFP57$^{-/-}$ dams (right). **Left -** WT pups raised by WT dams (black triangles) served as a baseline, compared to ZFP57$^{+/-p}$ pups raised by WT dams (light grey squares) or cross-fostered to WT dams (dark grey squares), and ZFP57$^{m-/+}$ (yellow circles) pups raised by or cross-fostered to WT dams. **Right -** WT pups raised by WT dams (black triangles) served as a baseline, compared to WT pups raised by ZFP57$^{-/-}$ dams (green triangles), ZFP57$^{m-/+}$ pups raised by ZFP57$^{-/-}$ dams (light purple circles) or cross-fostered to ZFP57$^{-/-}$ dams (dark purple circles), and ZFP57$^{m-/+}$ (blue squares) pups raised by or cross-fostered to ZFP57$^{-/-}$ dams. (*n*=24-32/group). **(C)** Female pup weights at weaning in the groups described in A (*n*=24-32/group). Data are mean ± SEM. One-way ANOVA with Tukey's corrections. **(D-F)** TD-NMR measurements show body compositions of female ZFP57$^{+/-p}$ and ZFP57$^{m-/+}$ offspring, including, fat mass percentage **(D)** lean mass percentages **(E)**, and lean/fat ratio **(F)**. *n*=8-15 pups/group from 2-3 litters. Data are mean ± SEM. One-way ANOVA with Tukey's corrections. **(G)** Fasting glucose of ZFP57$^{+/-p}$ and ZFP57$^{m-/+}$ female offspring. *n*=8-15 pups/group from 2-3 litters. Data are mean ± SEM. One-way ANOVA with Tukey's corrections. **(H)** Intraperitoneal glucose tolerance test (IP-GTT) of female ZFP57$^{+/-p}$ offspring (left) and ZFP57$^{m-/+}$ offspring (right). ZFP57$^{+/-p}$ offspring born to WT dams is plotted twice for comparison (light grey). *n*=8-15 pups/group from 2-3 litters. Data are mean ± SEM. **(I)** Area under the curve quantifying the performance of different groups of females in the IP-GTT in Figure 6D. *n*=8-15 pups/group from 2-3 litters. Data are mean ± SEM.

**Supplementary table 1 – Primers used in this study.**

| Gene | Forward primer | Reverse Primer |
|---|---|---|
| Magel2 | AGAGCGCATGTTCATTGGTG | CCTCTACGCAGGCATAAGGAT |
| Phlda2 | CGACGAGATCCTTTGCGAGG | GTGGAAAAACAGCTCCTTGGC |
| Peg3 | GAGTCCAGCTTGCCGAAGAT | ATCGGCTTGTCACTCACCTC |
| GnasXL | AGAACCTTTGGAAGCCCCAG | GTTCGGTCGCCATTTCTTCG |
| IGFII | ATCGTGGAAGAGTGCTGCTT | GTAGACACGTCCCTCTCGG |
| Mest | CCAAAAGCTCCTCAAAGACG | ACGGTCCAAAGACTGGAGTG |
| β Actin | GGCTGTATTCCCCTCCATCG | CCAGTTGGTAACAATGCCATGT |
| Grb10 | AAACAGGACCACCTCCCTCT | GCGGTTTGAATTGTCAAGGT |
| GATA3 | CTCGGCCATTCGTACATGGAA | GGATACCTCTGCACCGTAGC |
| Stat5a | CGCCAGATGCAAGTGTTGTAT | TCCTGGGGATTATCCAAGTCAAT |
| Stat5b | CGATGCCCTTCACCAGATG | AGCTGGGTGGCCTTAATGTTC |
| Stat6 | CTCTGTGGGGCCTAATTTCCA | CATCTGAACCGACCAGGAACT |
| C/EBPb | TGATGCAATCCGGATCAA | CACGTGTGTTGCGTCAGTC |
| Notch3 | AGATCAATGAGTGTGCATCC | GCAGACTCCATGACTACAGG |
| Elf5 | ATGTTGGACTCCGTAACCCAT | GCAGGGTAGTAGTCTTCATTGCT |
| β-Tubulin | TTCAGCTGACCCACTCACTG | AGACAGGGTGGCATTGTAGG |
| β-Actin | GGCTGTATTCCCCTCCATCG | CCAGTTGGTAACAATGCCATGT |
| GAPDH | AAGGGCTCATGACCACAGTC | GGATGCAGGGATGATGTTCT |
| Zac1 | TTCGTCACCCTGGAGAAGTT | GGTCTGGAGGTGGTTCTTCA |
| Rasgrf1 | TGATCGTATCCAATCCAGCA | ACACCACCTGGTTCCTCTTG |
| Snrpn | TAAATCTCAGCCCTTCTCTTCCC | AATGCAGTAAGAGGGGTCAAAAA |
| Nnat | CACCCACTTTCGGAACCAT | GCAGGGAGTACCTGAACACCT |
| IGF2R | AGCTAAATGGTGGCTATCTGGT | GGGTCGGCCAACGTCAAAT |
| Nespas | ACTGATCCTCTCGTCTGGGA | TCCGCGCAACTTTATAGGGC |
